# DNA Flipping as Facile Mechanism for Transmembrane Signaling in Synthetic Cells

**DOI:** 10.1101/2025.05.31.656672

**Authors:** Lorena Baranda Pellejero, Maria Vonk-de Roy, Doruk Baykal, Livia Lehmann, Andreas Walther

## Abstract

Transmembrane signaling is essential for cellular communication, yet reconstituting such mechanisms in synthetic systems remains challenging. Here, we report a simple and robust DNA-based mechanism for transmembrane signaling in synthetic cells using cholesterol-modified single-stranded DNA (Chol-ssDNA). We discovered that anchored Chol-ssDNA spontaneously flips across the membrane of giant unilamellar lipid vesicles (GUVs) in a nucleation-driven, defect-mediated process. This flipping enables internal signal processing through hybridization with encapsulated complementary DNA and activation of downstream processes such as RNA transcription. The phenomenon shows a high transduction efficiency, is generic across DNA sequences and lipid compositions, and can be enhanced by glycerol, which modulates membrane dynamics. Mechanistic insights using fluorescence microscopy, nuclease degradation assays, and membrane permeability assays reveal that flipping is dominated by transient membrane pores. Leveraging this facile translocation process, we demonstrate selective transcriptional activation inside synthetic cells, underscoring the potential of Chol-ssDNA flipping as a programmable tool for synthetic biology and bottom-up synthetic cell design.

## Introduction

Transmembrane signaling is a fundamental process that enables cells to communicate and respond to their environment. A variety of transmembrane proteins detect extracellular signals and initiate intracellular responses through signal transduction across the membrane. Receptor tyrosine kinases^1^ (RTKs) and G-protein-coupled receptors^2,3^ (GPCRs) are examples of signal transduction systems that operate without molecular matter exchange across the membrane. RTKs initiate signaling through ligand-induced dimerization, while GPCRs undergo global conformational changes upon ligand binding. In contrast, ion channels and pore-forming proteins create regulated openings in the membrane, allowing the selective passage of ions or small molecules, followed by activation of intracellular processes. Beyond proteins, lipids also contribute to signaling by organizing into lipid rafts, which can serve as platforms for protein clustering and signal transduction.^4^ Additionally, lipid asymmetry and uneven protein localization within bilayer leaflets can contribute to membrane curvature and tension, which in turn regulate signaling pathways.^5,6^ In this context, cell-penetrating peptides represent a unique class of molecules capable of interacting and translocating across membranes.^7^ Their translocation mechanism is believed to involve the formation of transient pores, allowing passage without causing extensive membrane disruption.^8,9^ Unlike antimicrobial peptides that induce cell death by forming stable pores,^10^ cell penetrating peptides maintain membrane integrity, making them valuable for therapeutic applications such as drug and nucleic acid delivery across the membrane.^11^

This ability of biological systems to transduce signals across membranes has attracted interest from various disciplines, including bioengineering, chemical nanosciences, synthetic biology, and synthetic cell research, to both replicate natural mechanisms and pioneer novel strategies for transmembrane communication. Recreating these processes not only deepens our knowledge of membrane dynamics but also facilitates the engineering of synthetic cells (SCs) able to respond to environmental cues.^12,13^ One synthetic approach involves designing amphiphilic molecules that embed within the membrane and undergo reactions upon recognition to signals, altering their hydrophilicity and exposing functional sites on the inner leaflet, thereby triggering downstream processes.^14–16^ Such systems often require complex synthesis and lack modularity. DNA nanotechnology, which exploits the predictable interactions and programmable self-assembly of DNA, has emerged as a powerful tool for engineering transmembrane signaling systems.^17,18^ Efforts have focused on designing DNA nanopores able to span membranes when functionalized with hydrophobic anchors.^19^ Some nanopores feature a lid mechanism, allowing gated openings controlled by specific signals.^20^ However, these designs are primarily focused on transporting small molecules or ions across membranes,^20^ limiting their ability to detect larger or more complex signals. To mimic natural receptors like RTKs and GPCRs, DNA-based constructs with hydrophobic anchors in the internal backbone have been shown to enable membrane insertion, and communication between the interior and exterior by externally triggered dimerization.^21–26^ Signal amplification is often necessary to render these processes measurable, and is typically achieved using catalytic DNA motifs, such as G-quadruplexes or DNAzymes, to enhance downstream reactions with detectable fluorescence signals.

Despite advances in DNA-based signal transduction, challenges remain, as these multimerization mechanisms are limited by the 2D diffusion of DNA receptors on the membrane and their ability to find each other for dimerization. Additionally, DNA can adopt multiple conformations on the membrane including anchored, non-directional, or partially embedded, rather than a strict transmembrane orientation.^22^ Recent efforts have shown an asymmetric DNA triplex that inserts directionally in a transmembrane manner,^24^ but such transmembrane DNA-based designs rely on multiple hydrophobic anchors with complicated synthesis, high costs, and low modularity.

In this study, we introduce extremely facile and spontaneous transmembrane information transfer, wherein commercial, cholesterol-modified single-stranded DNA (Chol-ssDNA) spontaneously translocates across lipid vesicle membranes, enabling transmembrane signaling and activating downstream processes inside such lipid-based SCs. We demonstrate that Chol-ssDNA anchors to the outer membrane leaflet and efficiently flips to the inner lumen. The process starts from a localized point and spreads across the entire liposome, suggesting a defect-nucleated process. This results in a high concentration of Chol-ssDNA in the inner leaflet. We evaluate this process by recruiting fluorescently tagged ssDNA encapsulated within the liposomes to the inner leaflet of the membrane. We show that the process is generic to many ssDNA strands, lipid compositions, and can be facilitated by glycerol altering membrane dynamics. We utilize this DNA flipping phenomenon as a transmembrane signaling process to control transcriptional activity within SCs. We anticipate this facile mechanism to contribute to a versatile tool for engineering SC communities and interfacing to biological systems.

## Results and Discussion

### DNA flipping across the membrane

The artificial signal transduction mechanism is built from three main components: a synthetic cell lipidic membrane, an external trigger in the form of a cholesterol-modified DNA strand (Chol-ssDNA), and encapsulated DNA-based circuits. Upon transmembrane signaling initiated by adding the Chol-ssDNA, followed by DNA flipping from the outer to the inner leaflet, the encapsulated circuit processes information within the SC interior and generates an output (Figure 1). This output can be engineered to trigger distinct downstream responses, such as inducing membrane fluorescence or initiating RNA transcription.

**Figure 1.**
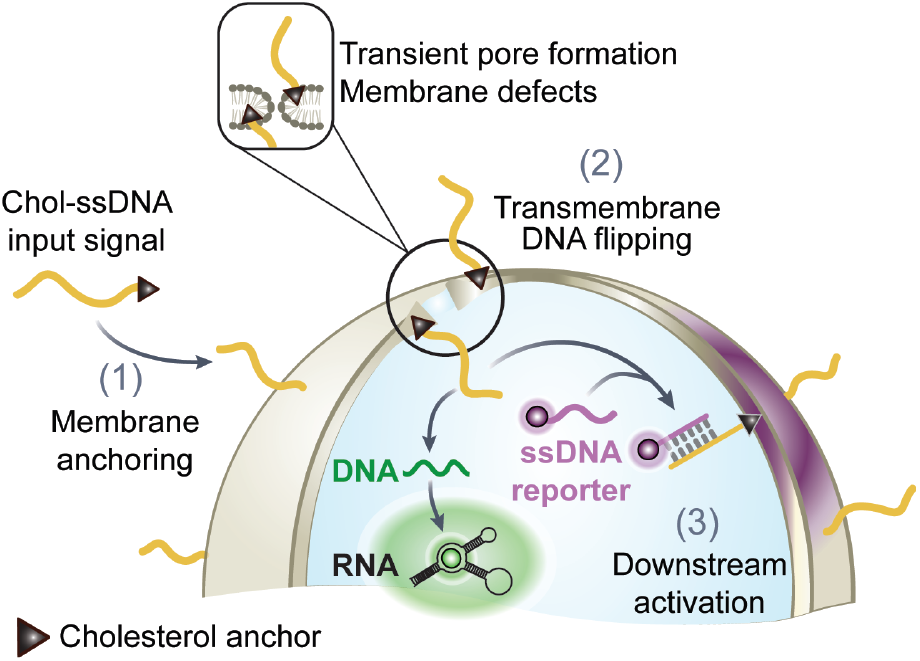
DNA Flipping as a Mechanism for Transmembrane Signaling in SCs. Cholesterol-modi.ied single stranded DNA (Chol-ssDNA) anchors to the outer lea.let of lipid vesicles. Upon pore formation and translocation, the Chol-ssDNA is exposed to the interior of the vesicle where it interacts with encapsulated DNA strands, leading to a membrane .luorescent signal by binding to a ssDNA reporter and simultaneously activating the transcription of RNA.

In more detail, to characterize the translocation mechanism of Chol-ssDNA, we prepared giant unilamellar vesicles (GUVs) composed of DOPC via a slightly modified emulsion transfer method^27,28^ and encapsulated a complementary dye-labeled ssDNA (Figure 2a). Confocal laser scanning microscopy (CLSM) visualizes this ssDNA reporter distributed homogenously in all GUVs (Figure 2b, left). To initiate the signal transmission, we added a 26-nt Chol-ssDNA (nt = nucleotide; modified at the 5’ end) having a 12-nt domain complementary to the encapsulated ssDNA reporter at the 3’ end. Chol-ssDNA anchors immediately to the outer membrane leaflet of all the vesicles (Figure S1). After a few seconds, CLSM reveals localization of the encapsulated ssDNA reporter from the cavity to the membrane in some GUVs, resulting in a dark cavity and bright fluorescence ring at the GUV membrane (Figure 2b, right image and zoom-in). This suggests that Chol-ssDNA translocates to the inner leaflet, where it binds the encapsulated ssDNA reporter. While flipping of cholesterol across lipid bilayers is a well-established phenomenon,^29^ critically, the translocation of Chol-ssDNA has not been previously reported. Despite multiple studies employing anchored Chol-ssDNA for various applications,^30^ this flipping behavior went unrecognized, likely because previous work never employed the appropriate experimental setup to monitor this phenomenon.

**Figure 2.**
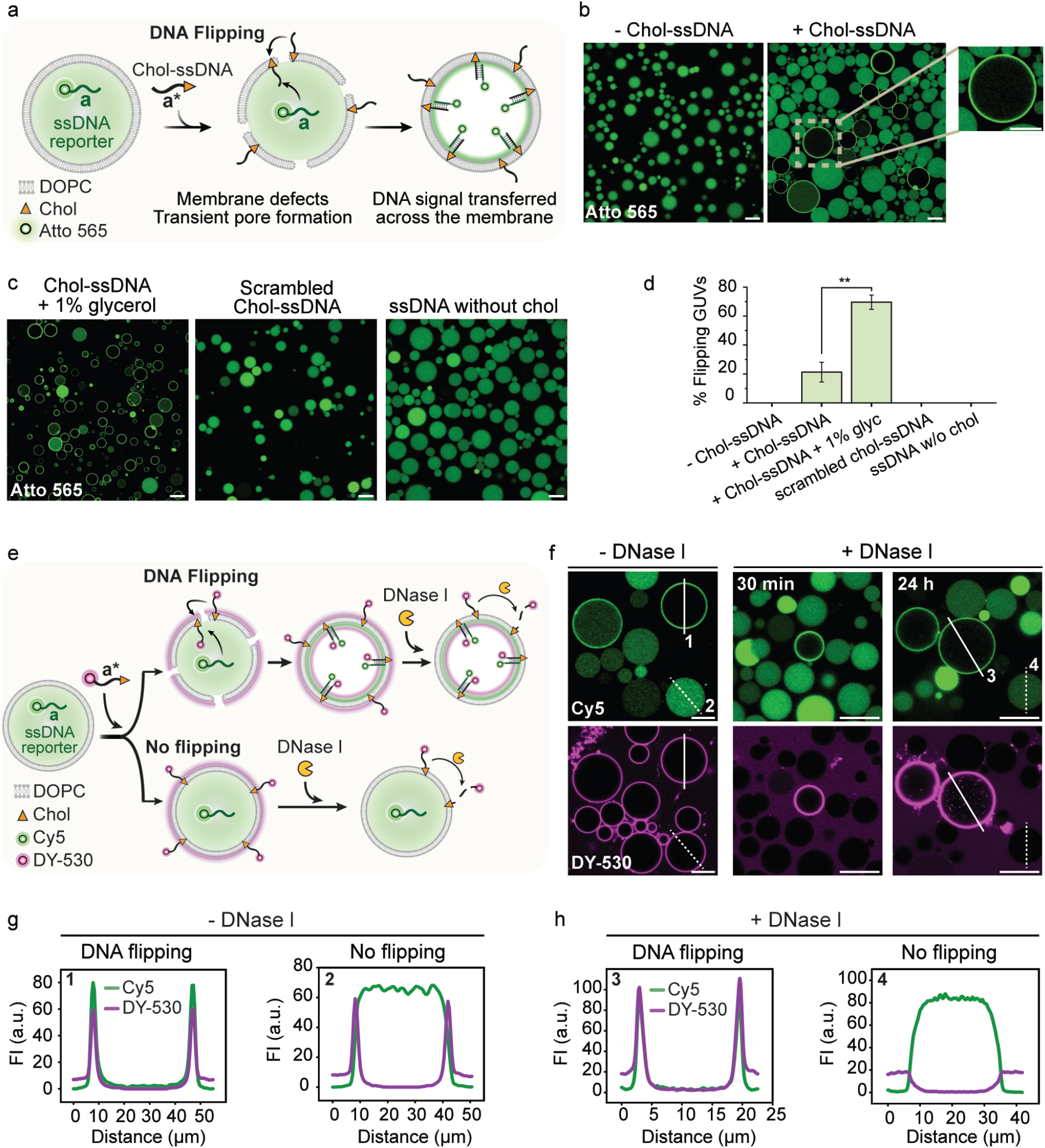
Flipping of end-cholesterol-modified single stranded DNA (Chol-ssDNA) across GUV membranes. **a** Schematic illustration of the translocation of an anchored Chol-ssDNA to the inner leaflet, facilitated by transient pores or defects in the membrane. The flipping of DNA is evidenced by hybridization with a Atto565-labeled ssDNA reporter encapsulated within the GUV. **b** CLSM image of GUVs with fluorescent ssDNA reporter encapsulated in absence and presence of Chol-ssDNA in the outer solution. Zoom-in on a GUV where DNA flipping occurred, with fluorescence localized at the membrane and depleted in the interior (right). **c** CLSM images show enhanced DNA flipping in GUVs with 1% v/v glycerol added to the outer solution (left) and under different control conditions, such as using scrambled Chol-ssDNA lacking strand complementarity (middle), and ssDNA signals without cholesterol modification (right). **d** Quantification of GUV flipping efficiency. The graph shows the percentage of flipping events in GUVs under the different conditions in panel b and c. **e** Schematic illustration of the nuclease degradation assay, depicting two possible pathways: (top) GUVs where flipped Chol-ssDNA remain protected from DNase I degradation; (bottom) GUVs without flipping, leaving Chol-ssDNA exposed on the outer membrane for complete degradation by DNase I. **f** CLSM images of GUVs before and after 30 min and 24 h of DNase I treatment. **g** Line profile analysis of selected regions of interest (ROIs) from CLSM images in panel f. The profiles belong to GUVs that did and did not undergo DNA flipping, before DNase I treatment and **h** after DNase I treatment. Data (d) are presented as the mean ± standard deviation (s.d.), with error bars representing the standard deviation from three independent experiments. Two-sided Student’s t-tests were conducted to assess statistical significance. Conditions: DOPC GUVs. Outer solution: 1xTE buffer (10 mM Tris, 1mM EDTA) pH 8, 150 mM NaCl, 400 mM glucose; 0.3 μM ssDNA scavenger (a*) (see Figure 3h) (panel a-d); 0.5 μM Chol-ssDNA (a*) (panel b right and panel c left); 1% v/v glycerol (panel c, left); 0.5 μM scrambled Chol-ssDNA: (panel c, middle); 0.5 μM ssDNA without chol (panel c, right); 0.3 μM ssDNA scavenger (b*) (panel e-h); 0.5 μM DY-530-labeled Chol-ssDNA (b*) (panel e-h); 1 g/L DNase I (panel e-h). Inner solution: 1xTE buffer pH 8, 150 mM NaCl, 400 mM sucrose; 1 μM Atto565-labeled ssDNA reporter (a) (panel a-d); 1 μM Cy5-labeled ssDNA reporter (b) (panel e-h). Scale bar: 20 µm.

This flipping process does not occur in all GUVs, but in a significant 20% of the sample (Figure 2d). It is common for liposomes to exhibit heterogeneous behavior in a population, although the underlying reasons for this variability are rarely studied.^31^ We hypothesized that fluidity of the membrane may play an important role and thus added 1% v/v glycerol to the outside solution after Chol-ssDNA addition to dynamize the membrane.^32^ Indeed, this increases the yield of transmembrane positives up to 70% (Figure 2c, left, Figure 2d). We discuss the effect of glycerol on DNA flipping further below. As a control, the addition of a scrambled Chol-ssDNA unable to hybridize to the encapsulated ssDNA reporter does not produce any accumulation of a fluorescent ring at the inner leaflet, confirming that the partitioning of the encapsulated ssDNA reporter from the cavity to the membrane is caused by sequence-specific hybridization (Figure 2c, middle). Moreover, adding non-modified ssDNA without cholesterol does not induce flipping either, highlighting the necessity of initial membrane anchoring for the translocation process to occur (Figure 2c, right).

The process is also independent of additives often added during imaging of GUVs. For instance, replacing polyvinyl alcohol (PVA) with BSA as an agent to passivate the glass surface during imaging still renders efficient flipping. PVA is known to interact and change the membrane properties (Figure S2).^33^ Additionally, DNA flipping also occurs in GUVs produced via a hydration protocol,^34^ thus avoiding any potential residual oil traces from the emulsion transfer method (Figure S3). Moreover, the effect of cations such as Na^+^ and Mg^2+^ on DNA flipping is minimal, although a minimum salinity (e.g., 150 mM NaCl or 2 mM MgCl_2_) is required for Chol-ssDNA to anchor to the membrane.^35^ These controls highlight that the mechanism is generic and independent of the GUV preparation method or imaging setup.

We employed a nuclease digestion assay to confirm that Chol-ssDNA translocates to the inner leaflet of the membrane and to track its location over time. In this assay, we used a 33-nt Chol-ssDNA with an additional DY-530 fluorophore at the 5’ end in combination with GUVs. Addition of DNase I can only degrade the Chol-ssDNA-DY-530 strand from the outside and would leave flipped constructs in the inside unharmed (Figure 2e). This assay also assesses whether Chol-ssDNA flips back to the outer leaflet over time, which would be indicated by a continuous and slow loss of membrane fluorescence. Experimentally, upon adding Chol-ssDNA-DY-530 to DOPC GUVs, the appearing membrane fluorescence confirms anchoring to all available GUV membranes (Figure 2f, left, DY-530 channel). GUVs where Chol-ssDNA-DY-530 translocates inward appear with bright rings and dark cavities in the green channel (Cy5) due to hybridization with the encapsulated ssDNA reporter (Figure 2f, Cy5 channel). After 30 minutes of DNase I digestion, DY-530 fluorescent rings corresponding to the flipped construct remain visible, whereas GUVs without flipping show complete loss of fluorescence (Figure 2f, middle). Even after 24 hours, fluorescence persists, indicating that the flipped Chol-ssDNA remains largely encapsulated within the GUV and flipping back to the outer leaflet would be a very slow process (Figure 2f, right).

To determine whether the flipping process is driven by hybridization with the encapsulated complementary ssDNA reporter, we repeated the experiment with an encapsulated scrambled ssDNA and for empty GUVs. Figure S4 shows similar results with digestion-resistant fluorescent rings. This confirms that flipping does not depend on the presence of a complementary ssDNA reporter.

### DNA flipping follows a nucleation-like defect-driven mechanism

To understand more mechanistic details of the process, we recorded timelapse CLSM data. Realtime imaging reveals that the translocation occurs in a rapid, nucleated, and concerted transition starting from one point of the GUV rather than gradually or through multiple stages. Typically, within tens of seconds, the anchored Chol-ssDNA molecules undergo complete flipping, shifting the GUV from one state to another (Figure 3a, b; Figure S5 for more flipping events). Strikingly, the bright fluorescence ring in the membrane appears asymmetrically, emerging from a specific point and rapidly spreading across the bilayer (Figure 3a, b; Supplementary Video 1). This localized initiation suggests that translocation is a defect-mediated process, where transient membrane disturbances create openings that allow anchored DNA to flip at significant amounts. This translocated DNA then diffuses across the entire GUV. Notably, in some instances DNA flipping initiates immediately after the bursting of an adjacent GUV, which generates disturbances in the surrounding environment (Supplementary Video 2). This suggests that any type of disturbance (such as nearby vesicle bursting or random vesicle collisions) may induce membrane defects that trigger DNA flipping. The depletion of the ssDNA reporter from the cavity happens simultaneously with the formation of a fluorescent ring, confirming that these two processes are connected (Figure 3c). Note that the diffusion of the Chol-ssDNA anchored on the membrane, as assessed by fluorescence recovery after photobleaching (FRAP, Figure S5) is in the order of 0.3 µm^2^/s and thus faster than the spreading of the fluorescent ring that requires several tens of seconds. This confirms that membrane constraints, such as small pores or defects, limit the spread of the fluorescent ring during flipping, not the diffusion rate of the Chol-ssDNA itself. Besides, the complete depletion of the complementary ssDNA from the cavity suggests that the amount of Chol-ssDNA undergoing flipping is high, as it binds all the available encapsulated ssDNA reporter. A theoretical calculation of the minimum amount of Chol-ssDNA that has flipped to the interior yields that at least 1,807 molecules per µm^2^ of the inner leaflet of the membrane, based on an average GUV diameter of 18 µm (Supplementary Note 1). This data corresponds to a distance of ∼ 15 nm between flipped Chol-ssDNA molecules, with one flipped Chol-ssDNA molecule present for ∼ 790 DOPC molecules.

**Figure 3.**
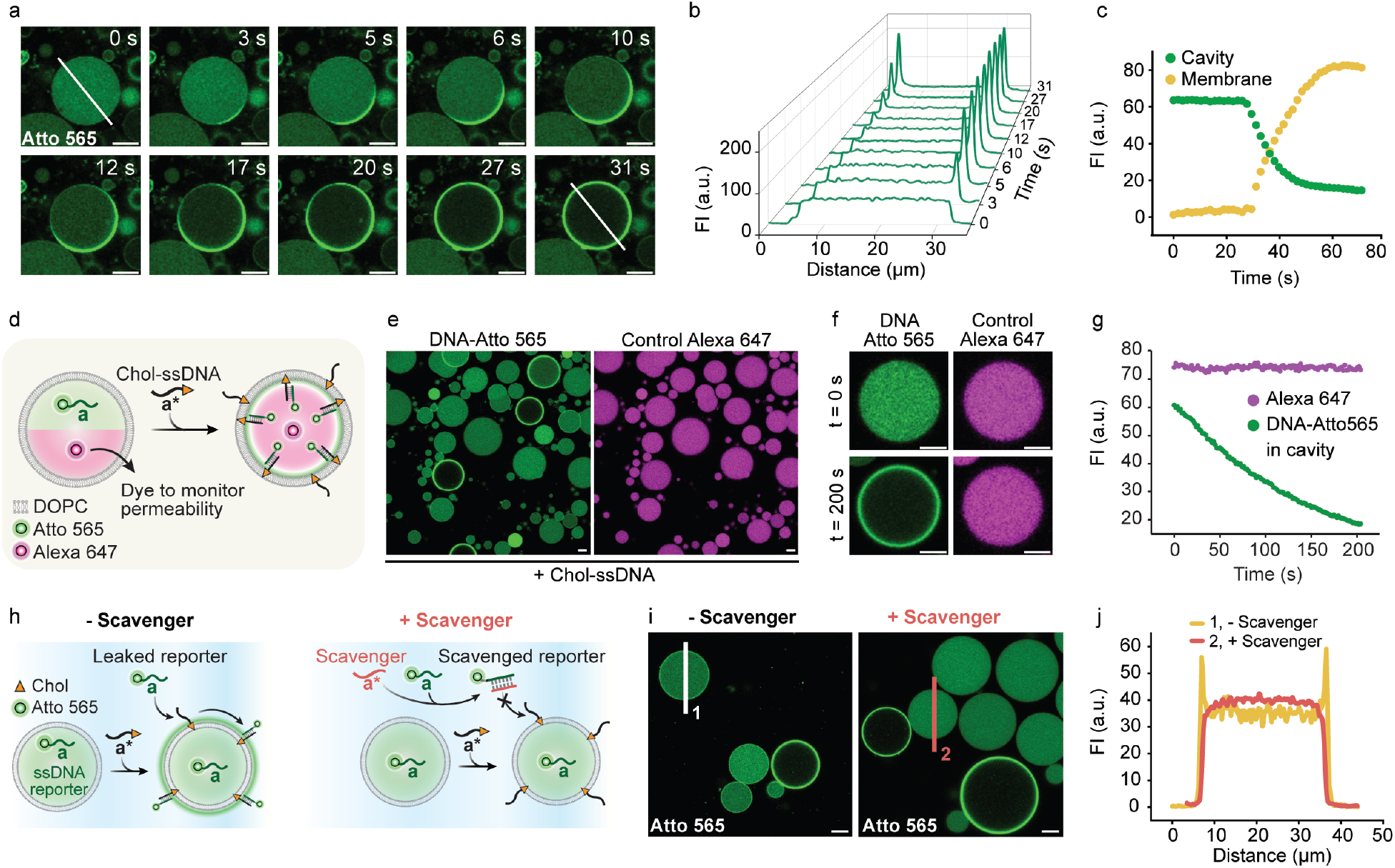
Real-time dynamics of Chol-ssDNA flipping provide evidence for a nucleation-driven mechanism. **a** Time-lapse CLSM images showing the flipping of Chol-ssDNA across a GUV membrane. The process is monitored through binding of fluorescent ssDNA reporter that hybridizes to the inner leaflet of the GUV membrane once the Chol-ssDNA flips to the inside. **b** Line profile analysis of DNA translocation over time according to the line in a. **c** Graph showing fluorescence intensity (FI) over time for the cavity and the entire membrane. **d** Schematic representation of the flipping experiment, in which Alexa 647 is co-encapsulated in the GUVs to monitor membrane permeability. **e** CLSM images of GUVs showing encapsulated DNA-Atto 565 reporter fluorescence (left) and Alexa 647 as a control for permeability (right). GUVs where flipping occurs are visible as rings in the DNA-Atto 565 channel and retain Alexa 647 within their lumen, indicating no leakage. **f** Time-lapse CLSM images of the same GUV before and after flipping (0 and 200 s) and **g** FI over time. **h** ssDNA reporter strands (a) that leak during GUV preparation must be scavenged by a scavenger ssDNA (a*) before addition of Chol-ssDNA (a*) signals to prevent false positives. **i** CLSM images showing GUVs with Chol-ssDNA, with and without the previous addition of scavenger ssDNA. In absence of scavenger, GUVs that did not undergo flipping show membrane fluorescence (false positives). Addition of scavenger fully prevents false positives. **j** Line profile analysis of the FI taken along the lines drawn in i. Conditions: DOPC GUVs. Outer solution: 1xTE buffer pH 8, 150 mM NaCl, 400 mM glucose; 0.3 µM ssDNA scavenger (a*); 0.5 μM Chol-ssDNA (a*). Inner solution: 1xTE buffer pH 8, 150 mM NaCl, 400 mM sucrose; 1 µM Atto565-labeled ssDNA reporter (a); 1 μM Alexa 647 (panel d-g). Scale bar: 10 µm.

In terms of the mechanism, we suggest that the translocation is induced by asymmetry and tension due to attachment of bulky Chol-ssDNA segments to the outside. The driving force for efficient translocation is rooted in equilibrating Chol-ssDNA concentration (inside and outside) and in releasing membrane tension on these large GUVs, because the anchoring of Chol-ssDNA exclusively on the outer membrane induces membrane curvature and perturbs the membrane.^36,37^ Changes in membrane curvature can be identified for GUVs that are assembled at one bilayer and where one of the GUVs undergoes flipping (Supplementary Video 3). Similar driving forces have been suggested for peptides.^8^

To investigate whether GUV leakage occurs during the flipping process and to rule out leakage of the encapsulated strand as the cause of the observed depletion, we co-encapsulated a moderately sized fluorescent molecule (Alexa 647, zwitterionic) with the fluorescent ssDNA reporter (Figure 3d). CLSM shows that Alexa 647 remains enclosed within the GUV while the depletion of the fluorescent strand happens. (Figure 3e-g). Hence, the flipping process does not involve the formation of large pores through which small molecules (∼900 Da) can pass.

One of the critical points is to avoid false positives — an issue that has thus far not been properly considered in the field of DNA transmembrane receptors. We found that the use of a scavenger ssDNA is crucial to prevent false positives (Figure 3h). During GUV preparation, minor leakage of the inner solution of a GUV transitioning through the emulsion interface in the emulsion transfer method is unavoidable. In plain solution, such low concentrations remain invisible in subsequent CLSM imaging due to dilution. However, upon adding Chol-ssDNA, its membrane localization enables hybridization with any leaked fluorescent and complementary ssDNA, concentrating it on the membrane and creating a peripheral ring that can be easily misinterpreted as transmembrane events (Figure 3i, left image). To eliminate this ambiguity, it is critical to introduce a complementary scavenger ssDNA into the outer solution post-GUV preparation and prior Chol-ssDNA signal addition. This scavenger binds and neutralizes any leaked fluorescent ssDNA reporter, ensuring no unintended membrane fluorescence (Figure 3i, j).

### Understanding the membrane structure during DNA flipping

For ssDNA to flip across the membrane, some form of membrane distortion is required, such as a transient toroidal pore. As previously mentioned, moderately sized molecules such as Alexa 647 (∼ 900 Da) remain encapsulated within the GUV during DNA flipping, suggesting that any defects or pores that might form in the membrane are likely to be small (Figure 3d-g). To further investigate membrane structure and permeability during DNA flipping, we performed several assays.

First, we tested whether ion passage occurs across the membrane during DNA flipping, as GUVs do not allow ion passage across the membrane unless a water pore is formed. To do this, we encapsulated an engineered protein able to sense Ca^2+^, named G-GECO,^38^ and added CaCl_2_ to the outer solution (Figure 4a). If Ca^2+^ can cross the membrane during DNA flipping, it will bind to the G-GECO protein and switch on its fluorescence. CLSM clearly shows that all GUVs, in which DNA flipping had occurred after adding Chol-ssDNA turn bright green due to the activation of the Ca^2+^ sensor. This confirms that ion-permeable membrane pores open during translocation, whereas no ion passage occurs in DNA-anchored GUVs (Figure 4b).

**Figure 4.**
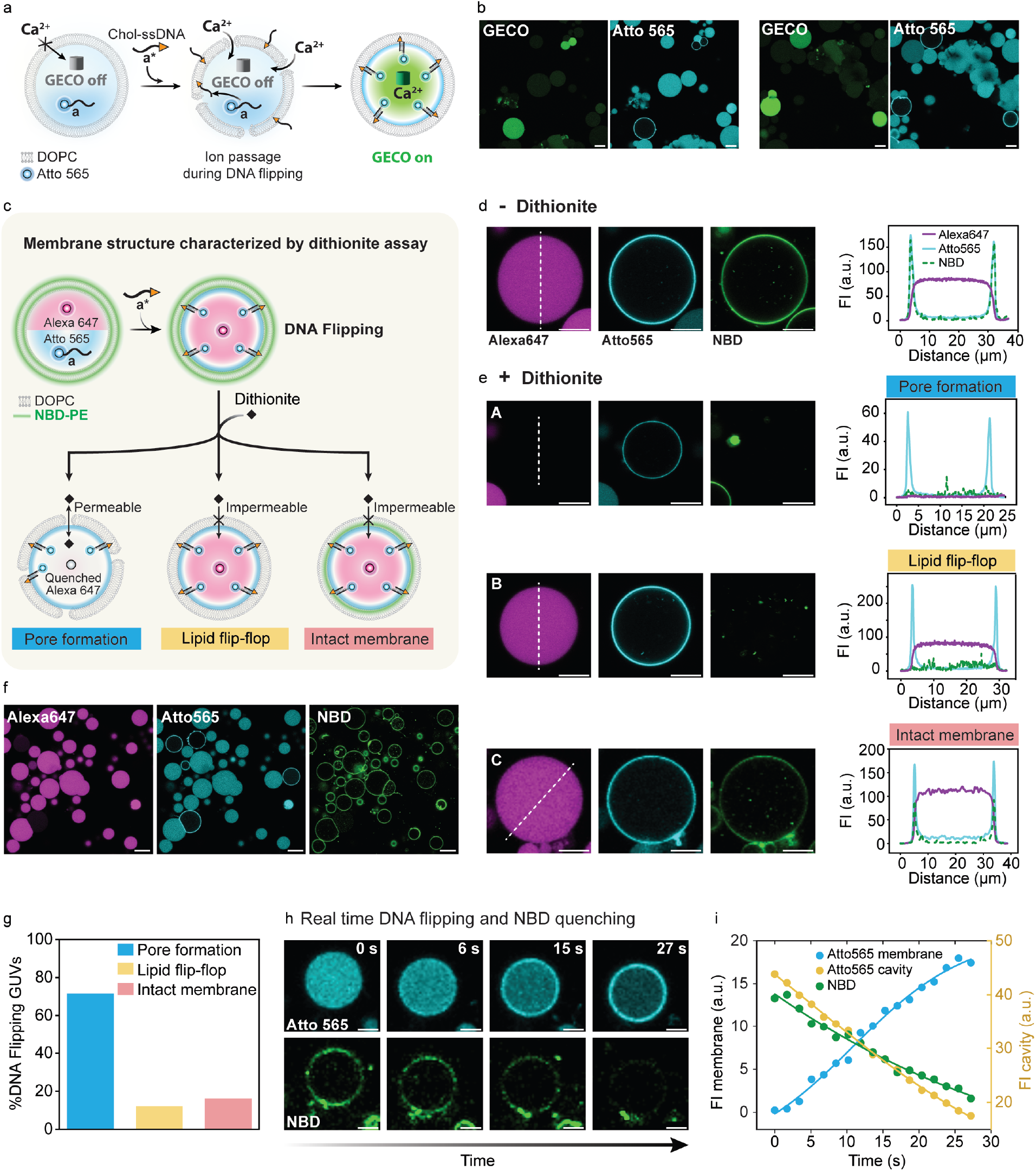
Ion permeability test and dithionite assay to probe the membrane structure of GUVs undergoing Chol-ssDNA translocation. **a** Ion permeability test using an encapsulated ion-sensitive protein sensor called G-GECO to assess whether Ca^2+^ can pass through the membrane while DNA flipping takes place. **b** CLSM images showing that DNA flipping is associated with G-GECO light up, confirming influx of Ca^2+^. **c** Schematic representation of the dithionite assay used to assess membrane integrity following Chol-ssDNA translocation. The assay evaluates membrane permeability to dianion dithionite, the occurrence of lipid flip-flop, or the preservation of an intact membrane. **d** CLSM images (left) and line profile analysis (right) of a GUV where DNA translocation occurred, before dithionite addition. **e** CLSM images (left) and line profile analysis (right) of GUVs after DNA translocation and dithionite addition reveal different membrane structures, highlighting sample heterogeneity. Some GUVs exhibit full quenching of both Alexa 647 and NBD, indicating membrane pore formation (Panel A). Others show complete NBD quenching while Alexa 647 fluorescence persists, suggesting lipid flip-flop without membrane permeabilization (Panel B). In contrast, some GUVs display NBD quenching only in the outer leaflet, with Alexa 647 fluorescence remaining, indicating an intact membrane (Panel C). **f** Overview CLSM image of the sample, showing a mix of GUVs with and without DNA translocation (DNA translocation monitored as rings in the Atto 565 channel). Most GUVs with flipped DNA exhibit pore formation, indicated by Alexa 647 and NBD quenching. **g** Quantification of membrane structure in GUVs after DNA translocation: A 72% of GUVs with flipped DNA exhibited pore formation, while 16% retained an intact membrane and 12% showed lipid flip-flop. **h** Time-series showing the Chol-ssDNA translocation process (Atto 565 channel) alongside the simultaneous quenching of NBD fluorescence (NBD channel). **i** Membrane FI over time for encapsulated Atto 565-ssDNA reporter localizing into the membrane and NBD membrane quenching, along with cavity FI to track Atto 565-ssDNA depletion. Line is a guide for the eye. Conditions: DOPC GUVs (panel a, b); DOPC + 0.1 mol% NBD-PE GUVs (panels c-i). Outer solution: 1xTE buffer pH 8, 150 mM NaCl, 400 mM glucose; 0.3 μM ssDNA scavenger (a*); 0.5 μM Chol-ssDNA (a*); 3 mM CaCl_2_ (panel a, b); 9 mM sodium dithionite (panel c-i). Inner solution: 1xTE buffer pH 8, 150 mM NaCl, 400 mM sucrose; 1 μM Atto565-labeled ssDNA reporter (a); 3 μM G-GECO (panel a, b); 1 μM Alexa 647 (panel c-i). Scale bar: 20 µm (panel b, f); 10 µm (panel d, e); 5 µm (panel h).

Second, we conducted a sodium dithionite quenching assay to assess membrane structure and lipid flip-flop, a rare process in plain GUVs.^39^ Dithionite is a membrane-impermeable reducing agent that quenches fluorescent NBD-labeled lipids (NBD = nitrobenzodiazole). To refine this assay, we encapsulated Alexa 647, a dye that is also quenched by dithionite,^40^ enabling us to detect pore formation and membrane rearrangements. Previous experiments confirmed that Alexa 647 remains enclosed within GUVs during DNA translocation (Figure 3d-g). Therefore, its fluorescence loss in this assay would indicate quenching due to dithionite entry via pores rather than GUV leakage. For this assay, we used GUVs labeled with NBD-PE and encapsulated Alexa 647 as well as a standard ssDNA reporter to track Chol-ssDNA flipping (Figure 4c). We first focus on GUVs with already occurred translocation events, as visible via the membrane ring in the Atto 565 channel (ssDNA reporter) and the NBD channel (labeled lipid), along with the cavity fluorescence from Alexa 647 (Figure 4d). After dithionite treatment, GUVs with flipped DNA display different populations (Figure 4f). Specifically, three distinct membrane behaviors can be identified. 72% of GUVs with flipped DNA exhibit complete quenching of both NBD and Alexa 647, indicating that the membrane remains permeable to dithionite even after DNA flipping (Figure 4e, panel A, Figure 4g). This may suggest the formation of very small water pores or defects, potentially stabilized by flipped DNA near these structures, as seen with certain peptides.^41^ Note that loss in Alexa 647 fluorescence does not come from leakage (see Figure 3d-g) but from ingress of dithionite (174 g/mol, dianion) quenching its fluorescence. 16% of GUVs with flipped DNA show partial NBD quenching (∼50%, corresponding to outer leaflet quenching) and no quenching of Alexa 647 (Figure 4e, panel C, Figure 4g). Pores for ingress of dithionite are absent and only the outer NBD lipids are quenched. A control experiment of NBD-GUVs without flipping shows a similar 50% NBD quenching (Figure S6). A minority of 12% of GUVs with flipped DNA retain Alexa 647 fluorescence while NBD is fully quenched. This can only arise from continuous lipid flip-flop behavior to quench all of it, while pores allowing ingress of dithionite are absent (Figure 4e, panel B, Figure 4g). In a further experiment, we added dithionite prior to Chol-ssDNA translocation to visualize in real time whether DNA flipping and NBD-lipid quenching occur simultaneously or whether one process precedes the other. Indeed, DNA insertion – evidenced by increased membrane fluorescence and cavity depletion in the Atto 565 channel – occurs simultaneously with the complete quenching of NBD-labeled lipids (Figure 4h, i). This confirms that both processes are connected.

Overall, this assay reveals that Chol-ssDNA translocation occurs via the formation of tiny pores, which are small enough to block dye passage yet large enough to allow anionic dithionite and Ca^2+^ ions to permeate. Following DNA flipping, the membrane structures show a slight heterogeneity among GUVs, with GUVs exhibiting pore formation, lipid flipping, or remaining intact. Most GUVs form ultrasmall pores able to transmit dithionite but not Alexa 647 that persist even after translocation. Whether these pores are dynamically reorganized (open and closing) or persistent is experimentally inaccessible to our knowledge.

### Influence of lipid composition, glycerol, and DNA sequence on DNA membrane translocation

The DNA flipping mechanism is driven by the anchoring of Chol-ssDNA to the lipid membrane, as well as membrane defects or changes in curvature. We hypothesized that variations in lipid composition, which influence fluidity, permeability, and lipid packing, may impact DNA flipping across the membrane. Initially, we characterized DNA flipping in DOPC GUVs, a zwitterionic lipid with a melting temperature (*T*_m_) of -17 °C,^42^ wherein ∼20% of GUVs exhibit DNA flipping (Figure 5a, b). Upon addition of 50 mol% cholesterol to increase membrane rigidity and reduce fluidity, the amount of GUVs undergoing DNA flipping decreases slightly to 15% (Figure 5a, b). This reduction aligns with our expectation that increased rigidity limits the membrane’s ability to facilitate DNA translocation. We also tested POPC, another zwitterionic lipid with a higher *T*_m_ of -2 °C and lower fluidity than DOPC.^42^ In POPC GUVs, 10% of GUVs exhibit DNA flipping, which is only half of the flipping efficiency observed in pure DOPC (20%), further highlighting the importance of lipid fluidity. Adding 50% cholesterol to POPC does not significantly affect flipping efficiency. Next, we examined Egg PC, a natural lipid mixture commonly used in the literature for transmembrane signaling in GUVs.^16,21,24^ Egg PC has an intermediate *T*_m_ of -15 °C compared to DOPC and POPC and is also neutral.^43^ Interestingly, Egg PC GUVs show the lowest DNA flipping efficiency, with only 4% of GUVs undergoing flipping, again also unaffected by the addition of 50 mol% cholesterol (Figure 5a, b). These findings suggest that factors beyond *T*_m_ influence DNA translocation, for instance, Egg PC might have an increased capacity to adapt to localized membrane stress due to its mix of lipids that can reorganize in response to local curvature stress. This reduces the necessity for translocation to relieve stress.

**Figure 5.**
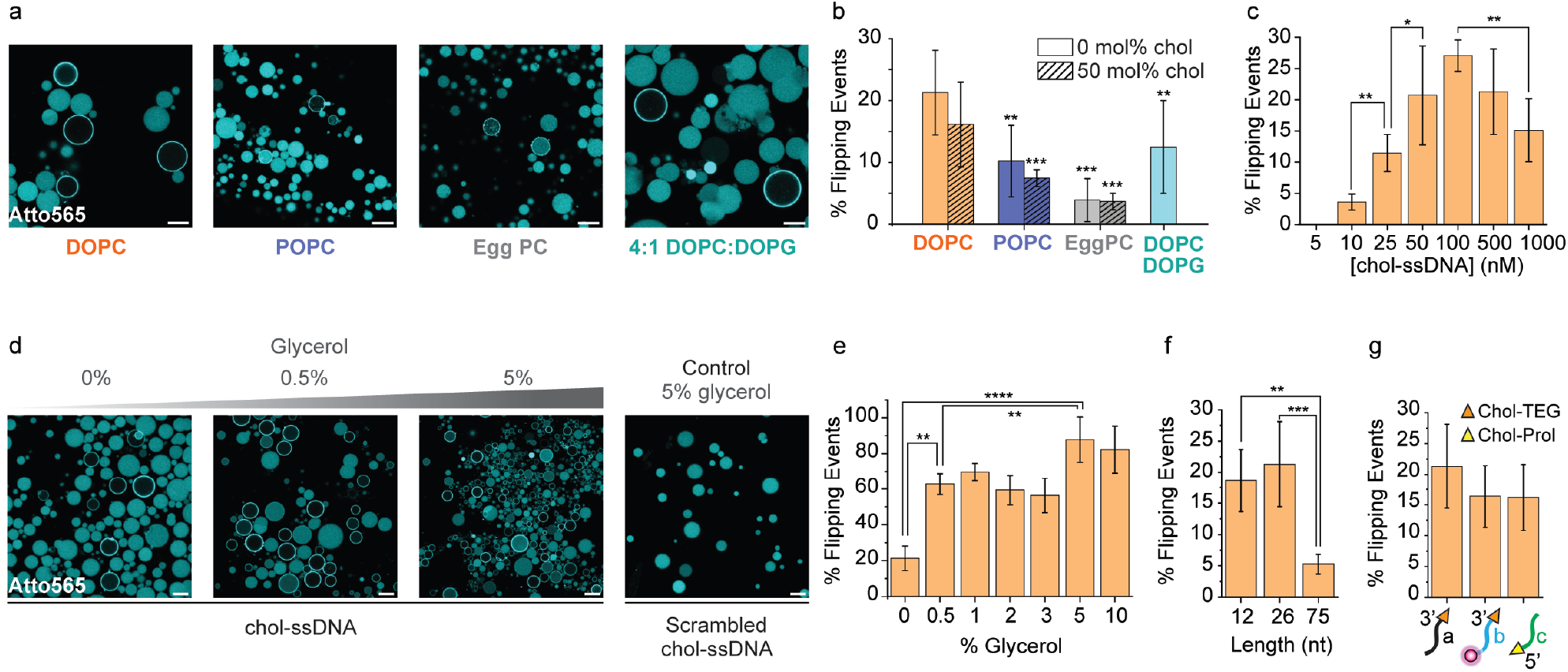
Effect of lipid composition, glycerol, Chol-ssDNA concentration and sequence, as well as anchor groups on the flipping process. **a** CLSM images and analysis showing how different lipid compositions and cholesterol content in the GUV influence the extent of Chol-ssDNA flipping. **b** Bar chart showing the percentage of GUVs undergoing flipping for different lipid compositions. **c** Titration of Chol-ssDNA showing how its concentration in solution affects the efficiency of flipping. **d** CLSM images depicting the effect of increasing glycerol concentration on Chol-ssDNA flipping. A control condition with 5% v/v glycerol and scrambled Chol-ssDNA is included to assess specificity. **e** Quantification of flipping events as a function of glycerol concentration. **f** Extent of DNA flipping using Chol-ssDNA of varying lengths. **g** Quantification of DNA flipping events using different Chol-ssDNA structures: sequence a (CholTEG at the 3’ end), sequence b (CholTEG at 3’ end and a DY-530 fluorophore at the 5’ end), and sequence c (cholesterol-prolinol at the 3’ end). Structure of CholTEG and Chol-prolinol can be found in Figure S7. Data are presented as the mean ± standard deviation (s.d.), with error bars representing the standard deviation from three independent experiments. Two-sided Student’s t-tests were conducted to assess statistical significance. Conditions: lipid as indicated in the figure (panel a, b); DOPC GUVs (panel c-g). Outer solution: 1xTE buffer pH 8, 150 mM NaCl, 400 mM glucose; 0.3 μM ssDNA scavenger (a*) (panel a-f); 0.5 μM Chol-ssDNA (a*) (panel a-e, unless otherwise specified in panel c); 0.5 μM scrambled Chol-ssDNA (panel d, right); 0.5 μM Chol-ssDNA (12-nt, 26-nt or 75-nt) (panel f); 0.5 μM Chol-ssDNA (sequence a, b or c) (panel g, see SI for corresponding scavenger sequences); Glycerol added at the concentrations specified in panel d and e (% v/v). Inner solution: 1xTE buffer pH 8, 150 mM NaCl, 400 mM sucrose; 1 μM ssDNA reporter (a*) (panel a-f); 1 μM ssDNA reporter (panel g, see SI for corresponding reporter sequences). Scale bar: 20 µm.

To investigate the role of membrane charge, we tested whether including a negatively charged lipid would hinder DNA translocation, given that DNA itself is negatively charged. We studied DNA translocation in 4:1 DOPC:DOPG GUVs, where DOPG is a negatively charged lipid, but detected no significant difference compared to zwitterionic DOPC GUVs (Figure 5a, b). This suggests that a negative membrane charge does not significantly inhibit DNA translocation, possibly because the cationic counter ions (Na^+^) in DNA and in the phospholipids screen electrostatic repulsion. Notably, the 4:1 DOPC:DOPG GUV system carries a negative charge, resembling a typical anionic plasma membrane.^44^ One may thus hypothesize that the observed DNA translocation mechanism may also be relevant in cellular environments.

Next, we examined the effect of varying Chol-ssDNA concentrations on DNA flipping. Concentrations below 10 nM do not induce flipping, indicating that a minimum threshold of anchored DNA is required to destabilize the membrane and enable translocation (Figure 5c). This is similar to peptide translocation mechanisms, where a critical concentration of adsorbed peptides is needed to nucleate a water pore.^41^ The highest flipping efficiency is achieved for 100 nM Chol-ssDNA. However, increasing the concentration to 1000 nM resulted in reduced DNA flipping, potentially due to increased membrane rigidity from the cholesterol anchor on DNA, as well as DNA crowding on the outer surface of the GUVs, which may hinder the flipping process.

Moreover, the addition of glycerol to the outer solution following the introduction of Chol-ssDNA significantly increases translocation events, with up to 80% of GUVs undergoing DNA flipping (Figure 5d, e). Even small amounts of glycerol, as low as 0.5% v/v, have drastic effects with increasing flipping events to 60% of the GUV population, while higher concentrations (5 to 10 %v/v) further increase the flipping efficiency up to 80% (Figure 5e). Glycerol is known to displace water molecules at the membrane surface by interacting with phospholipid heads, promoting lipid-lipid interactions and increasing lipid packing.^32^ Replacing glycerol with glucose in the GUV solution as an osmotic control did not enhance flipping (data not shown), suggesting that glycerol’s ability to increase flipping is not due to osmotic stress, but rather arises from glycerol–lipid interactions. When GUVs were pre-equilibrated with 1% (v/v) glycerol prior to the flipping experiment, the enhancement in flipping is absent, underscoring the relevance of an asymmetric initial distribution and membrane interaction to promote flipping. Additionally, fusion events occur upon adding glycerol, which are rarely seen in its absence (Supplementary Video 4). These results confirm strongly increased membrane dynamicity that facilitates pore nucleation and DNA translocation.

In addition to investigating the effects of membrane composition and membrane-interacting molecules like glycerol, we explored how variations in the DNA sequence itself can influence the flipping process. We expected the length of Chol-ssDNA to be a critical factor in its ability to undergo translocation, as long sequences may be bulky and too charged to efficiently cross the membrane. We thus tested sequences of different lengths (12 and 75-nt, in addition to the previously used 26-nt) while keeping the 12-nt complementary region to the encapsulated ssDNA reporter constant. The 12-nt Chol-ssDNA also undergoes DNA flipping with an efficiency similar to that of the previously studied 26-nt Chol-ssDNA. However, the 75-nt Chol-ssDNA shows a considerable drop in DNA flipping efficiency, with only 5% of GUVs exhibiting flipping events in absence of glycerol (Figure 5f). Additionally, we examined how the end group modifications of Chol-ssDNA affect flipping efficiency. The 26-mer Chol-ssDNA used throughout the study, containing a cholesterol-triethylene glycol (CholTEG) at the 3’ end, was compared to similar-length variants with an additional fluorophore (DY-530) added at the 5’ end and with a strand where CholTEG was replaced by cholesterol-prolinol at the 5’ end (structure in Figure S7). All sequences exhibit comparable flipping efficiencies (∼15%), with only a slight, non-significant increase observed in the original strand (20%) (Figure 5g). Therefore, the flipping mechanism remains effective for a broad range of DNA sequences and anchoring end groups, with a slight loss of translocation efficiency for longer DNA strands.

### Transmembrane signaling via DNA flipping activates RNA transcription

Having gained a deep understanding of the flipping process, we then focused on harnessing the DNA flipping mechanism to enable transmembrane communication between the external environment and the interior of the GUVs. Our goal was to encapsulate an internal processing circuit within the GUVs that can be triggered from the outside by transducing signals using the Chol-ssDNA flipping mechanism. By leveraging this approach, we aimed to activate a cell-free transcription system to produce RNA on demand—exclusively within the GUVs that undergo flipping.

To this end, we chose to transcribe a fluorescent RNA spinach aptamer.^45^ To achieve control over aptamer transcription via transmembrane signaling, we incorporated the T7 promoter sequence into a T7p*-Chol-ssDNA, and provided the template encoding for the RNA aptamer, as well as the T7 RNA polymerase, the NTP mix, and the DFHBI compound needed for the aptamer to light up inside the GUVs (Figure 6a). In addition to detecting RNA production, we monitored DNA flipping by co-encapsulating a Cy5.5-labeled ssDNA reporter, which provides fluorescent membrane feedback when flipping occurs (Figure 6a). In absence of the T7p*-Chol-ssDNA, the cavities of the GUVs show homogeneous fluorescence in the Cy5.5 channel, whereas the DFHBI channel remains dark. This confirms absence of RNA production (Figure 6b, top left). After adding T7p*-Chol-ssDNA and incubation for 3 h at room temperature, GUVs that underwent DNA flipping, as indicated by fluorescent membrane rings in the Cy5.5 channel, clearly display DFHBI fluorescence inside their cavities (Figure 6b, right). This demonstrates the activation of the RNA Spinach transcription. Around 80% of GUVs in which flipping occurred show RNA production, highlighting the link between membrane translocation and transcriptional activation. RNA production is completely absent in GUVs lacking DNA flipping. As a control, a scrambled Chol-ssDNA does not induce RNA transcription neither (Figure 6b, bottom left). The lack of RNA transcription in the remaining 20% of GUVs with flipped DNA may result from leakage of essential components (e.g. T7 RNA polymerase) during GUV preparation, a common cause of inter-vesicle variability, or from random permeability to nucleotides in a small fraction of GUVs, as observed in previous studies.^31,46^ Overall, this confirms that the DNA flipping technology is robust in the context of SC applications, because transcription requires specific buffers, enzymes, and also the nucleotides.

**Figure 6.**
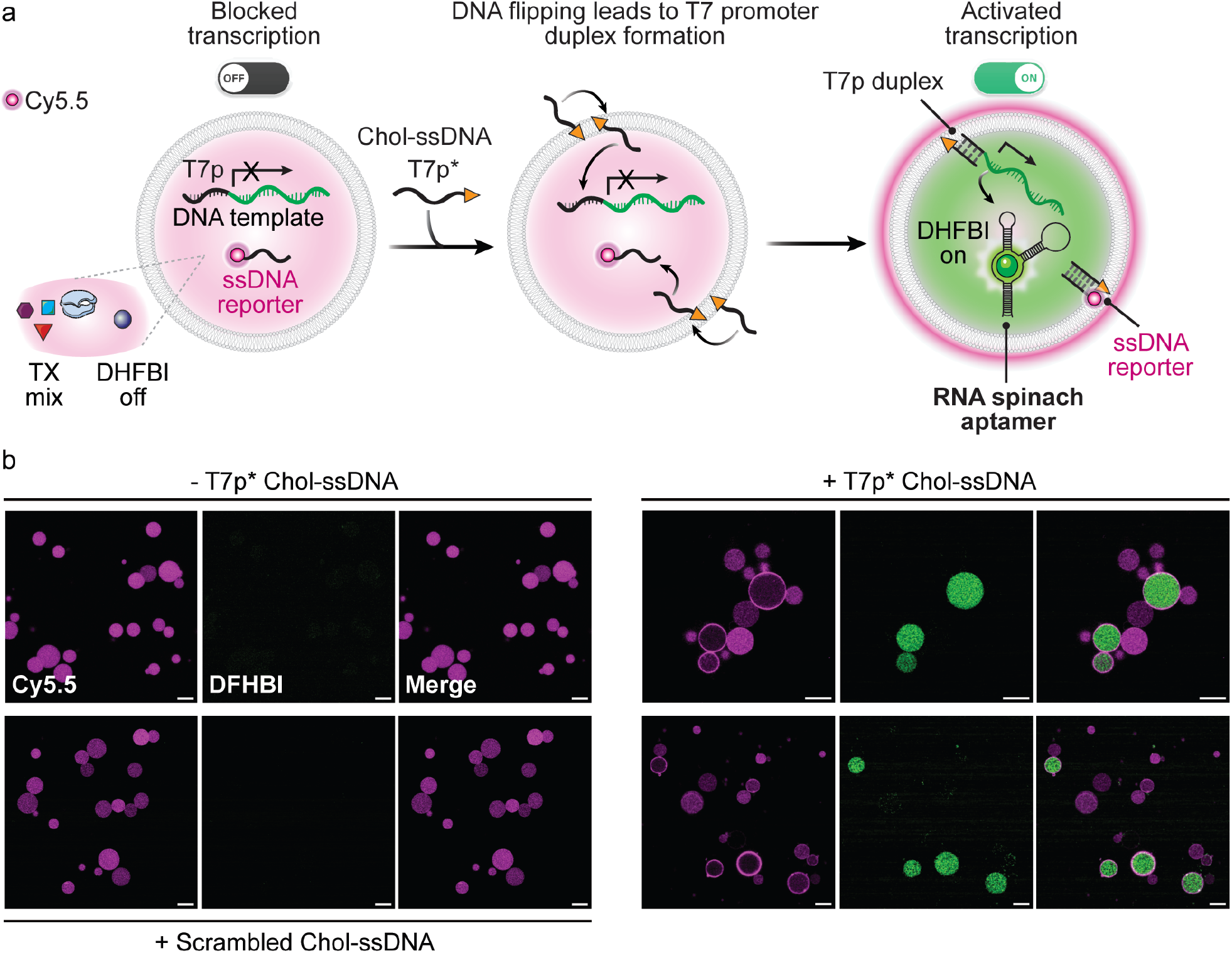
Transcriptional activation via DNA flipping in GUV-based SCs. **a** Schematic representation of the Chol-ssDNA flipping-enabled transcription in SCs. The cholesterol-modified T7 promoter sequence flips inward, allowing it to bind the DNA template and form the T7p duplex, which is essential to initiate RNA transcription. A fluorescent ssDNA reporter provides visual feedback on the flipping process. **b** CLSM images showing Cy5.5-labeled ssDNA reporter fluorescence (magenta) and DFHBI fluorescence (green) before and after incubation with T7p*-Chol-ssDNA. Following DNA flipping, DFHBI fluorescence appears inside flipped GUVs, confirming RNA Spinach aptamer production. GUVs without flipping show no transcriptional activation. GUVs incubated with scrambled Chol-ssDNA show no DFHBI fluorescence, confirming that transcriptional activation is sequence-specific (incubation at 25°C for 16 h). Conditions: DOPC GUVs. Outer solution: 40 mM HEPES, 125 mM KCl, 5 mM MgCl_2_, 400 mM glucose; 0.1 μM ssDNA scavenger (c*); 0.5 μM T7p*-Chol-ssDNA (where used); 0.5 μM scrambled Chol-ssDNA (where used). Inner solution: 40 mM HEPES, 125 mM KCl, 5 mM MgCl_2_, 400 mM sucrose; 6.7 mM each NTP; T7 RNA polymerase (added according to the kit instructions, see SI); 5 µM DFHBI; 1 µM DNA template; 0.3 µM Cy5.5-labeled ssDNA reporter (c). Scale bar: 20 µm.

## Conclusion

Our work has revealed a striking and previously unrecognized behavior of cholesterol-modified ssDNA: Beyond merely anchoring to lipid membranes — a well-established application in the field — these constructs can spontaneously translocate across the bilayer into the interior of GUVs. This challenges the assumption that Chol-ssDNA remains confined to the outer leaflet and, moreover, demonstrates a fundamentally new mode of transmembrane signaling for the engineering of SCs. The flipping process is initiated from a local membrane region and rapidly spreads across the entire GUV, consistent with a nucleation-driven mechanism involving membrane stress due to localized changes in curvature and transient pore formation.

We systematically dissected this phenomenon using real-time imaging, enzymatic degradation assays, and permeability tests, confirming the formation of tiny pores during translocation that are permeable to ions but not to dyes and which do not compromise overall membrane integrity. The flipping efficiency is highly robust and versatile. It can be observed for many different lipids, Chol-ssDNA of different length and sequence, as well as for different anchoring groups. Addition of glycerol to the GUVs largely boosts the transmembrane signaling up to 80% of the GUV population due to increasing membrane dynamics. Interestingly, negatively charged lipids still allow for translocation, offering a perspective for direct Chol-ssDNA translocation across cell membranes. We also identified the critical importance of scavenger strands to eliminate false positives, an important refinement for the field of synthetic DNA transmembrane receptors.

Beyond the implications for fundamental membrane biophysics, this work lays the groundwork for engineering programmable SCs. By coupling the flipping mechanism to a cell-free transcription system, we demonstrated a simple but versatile signal transduction route for activating internal processes in SCs in the context of RNA light-up aptamer transcription. This programmable signaling strategy eliminates the need for protein-based channels or GUV fusion assays, offering a minimalistic yet powerful alternative for engineering responsive synthetic cell systems. Looking ahead, integrating this DNA-flipping approach with more complex processing networks or applying it in biological membranes could pave the way for advanced SC communication, biosensing, and drug delivery platforms.

## Acknowledgments

We would like to thank Prof. Viktor Stein and Klara Eisenhauer for kindly providing the calcium protein sensor. L.B.P. acknowledges the support of the Alexander von Humboldt Foundation. L.B.P. and D.B. acknowledge support from the RTG 2516 “Structure Formation of Soft Matter at Interfaces”. This research was funded by the German Research Foundation (DFG) in the framework of the CRC 1552; Project No. 465145163. A.W. acknowledges funding from the Gutenberg Research Council Mainz underpinning his Life-Like Materials Program, the German Research Foundation grant WA 3084/19-1, the Max Planck Fellowship, and from the EU in the framework of the ERC Consolidator Grant to AW – M3ALI (101001638).

## Author information

### Associated Content

Supplementary Information. Materials and instrumentation, experimental details, image analysis methods, supplementary figures, theoretical calculations and video files.

## Table of Contents

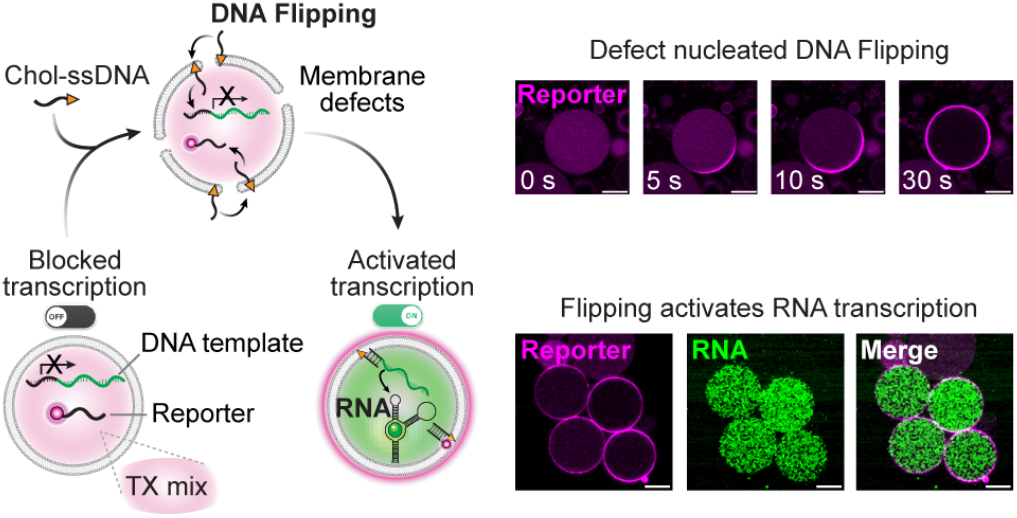

