## Supplementary Information for "DNA Flipping as Facile Mechanism for Transmembrane Signaling in Synthetic Cells"

### Supplementary Experimental Section

#### Materials

All chemicals were used without further purification and purchased from commercial sources. All DNA oligos were purchased HPLC-purified from Biomers. Supplementary Table 1 summarizes all sequences used in this study. Each DNA strand was dissolved in nuclease-free water and stored at -20 °C. Nuclease-free water, HiScribe T7 High Yield RNA Synthesis Kit including the NTP buffer mix (20 mM each NTP) and T7 RNA Polymerase mix were bought from New England BioLabs (NEB). DFHBI was purchased from Merck Millipore. DNase I (lyophilized), bovine serum albumin (BSA), cholesterol, hexadecane, sodium chloride, magnesium chloride, potassium chloride, calcium chloride, glucose, sucrose, sodium hydroxide, Tris base, HEPES, dithionite, polyvinyl alcohol (PVA, 99% hydrolyzed MW ≥ 100 kDa) and mineral oil were purchased (as bioreagent grade if available) from Sigma Aldrich. RNase-free 1xTE buffer pH 8 (containing 10 mM Tris adjusted to the specified pH with HCl and 1 mM EDTA), chloroform 99.9 % extra dry over molecular sieves and Alexa Fluor™ 647 Carboxylic Acid (referred to as Alexa 647 throughout this work) were purchased from ThermoFisher Scientific. DOPC (1,2-dioleoyl-sn-glycero-3-phosphocholine), POPC (1-palmitoyl-2-oleoyl-glycero-3-phosphocholine), Egg PC (egg yolk phosphatidylcholine; see below for the complete composition), DOPG (1,2-dioleoyl-sn-glycero-3-phospho-(1'-rac-glycerol) sodium salt) and 18:1 NBD-PE (1,2-dioleoyl-sn-glycero-3-phosphoethanolamine-N-(7-nitro-2-1,3-benzoxadiazol-4-yl) ammonium salt) were all purchased from Avanti Research. Egg PC is a natural lipid mixture composed of both saturated and unsaturated lipids, including: 0.2% of the 14:0 lipid, 32.7% 16:0 lipid, 12.3% 18:0 lipid, 1.1% 16:1 lipid, 32% 18:1 lipid, 17.1% 18:2 lipid, 0.2% 20:2 lipid, and 0.3% 20:3 lipid, 2.7% 20:4 lipid and 0.4% 22:6 lipid (Source: Avanti Research product description). Lipid stock solutions were prepared in chloroform and stored at -20 °C. Glass bottom 96-well imaging microtiter plates were purchased from VWR. The $Ca^{2+}$-responsive fluorescent protein sensor, G-GECO, was expressed and purified from E. coli.[1,2]

#### Instruments

Incubation was carried out on an Eppendorf ThermoMixer C with heated lid. DNA concentrations were determined by a DS-11 Spectrophotometer (DeNovix). Confocal laser scanning microscopy (CLSM) was performed on a Leica Stellaris 5 microscope with LasX software (v4.3.0.24308). Four laser lines and three HyD S detectors were used with a 63x objective (PL APO, oil immersion, 1.40 NA). Ultrapure water was obtained using a PURELAB Flex 3 system (ELGA).

#### Methods

##### GUV preparation by emulsion transfer

All aqueous solutions were made using ultrapure water. Inner solutions were prepared with 1xTE buffer pH 8 (containing 10 mM Tris, 1 mM EDTA), 150 mM NaCl, 400 mM sucrose and outer solutions contained 1xTE buffer pH 8, 150 mM NaCl, 400 mM glucose. 80 µL of a 25 g/L stock solution of chloroform-solubilized lipids (DOPC, unless otherwise specified), with or without 0.1 mol% of the fluorescent lipid 18:1 NBD-PE, was used to later prepare a lipid-in-oil solution at a final lipid concentration of 3 g/L in a total volume of 650 µL. To do this, a thin film of dried lipids was first formed by evaporating chloroform under a nitrogen flow in a 5-mL glass vial. Pre-dried mineral oil (vacuum-dried and stored over molecular sieves) was then added to the vial to a total volume of 650 µL. The mixture was sonicated at 55 °C for 45 minutes to enhance lipid solubilization in the oil, then allowed to cool down to room temperature. Following this, the lipid monolayer was prepared. To achieve this, 500 µL of outer solution is added to a 2-mL Eppendorf tube, followed by carefully layering 200 µL of the lipid-in-oil solution on top. The tube is then incubated at room temperature for 45 minutes, allowing an interfacial lipid monolayer to develop. The formation of the monolayer is indicated by interface flattening. Next, 450 µL of the lipid-in-oil solution and 7 µL of inner solution are combined in a 1.5 mL Eppendorf tube. The mixture is vortexed vigorously for 30 s or, if the inner solution contains proteins (main text Figure 6), pipetted up and down instead. This creates a water-in-oil emulsion, which is then transferred into the 2 mL Eppendorf tube containing the lipid monolayer interface. Immediately after, the tube is centrifuged at 4500 × g for 10 minutes at 5 °C. Following centrifugation, the upper oil phase is carefully removed, and the aqueous phase is gently pipetted up and down to redisperse the GUVs, forming the final GUV solution.

##### Surface treatment of the microtiter well plates

Proper surface treatment of the glass bottom wells of the 96-well plate is essential to prevent GUV adhesion and bursting. To achieve this, a 5% (w/w) PVA stock solution was prepared by dissolving the solid in ultrapure water under stirring at 90 °C. Then, 80 µL of a 1:10 dilution of the PVA stock solution (final concentration of 0.5% (w/w)) was added to each glass-bottom well, ensuring full coverage, and was completely removed immediately after. This process leaves a thin, evenly distributed coating on the glass surface. The microtiter plate was subsequently placed in an oven at 70 °C for 45 minutes and then allowed to cool to room temperature. The coating must be completed prior to the experiment.

In a control experiment, BSA replaced PVA as the coating agent (Figure S3). For this, 100 µL of a 10 g/L BSA solution was added to the well and incubated for 40 minutes. After incubation, the solution was removed, the well was washed once with 1xTE buffer pH 8, and the GUV solution was added.

#### Imaging by confocal laser scanning microscopy

The produced GUVs can be directly observed within the microtiter plate without any further sample preparation. A 50 µL volume of GUV solution was placed into the coated well and incubated for 2 min for the GUVs to settle to the bottom due to the density gradient between the intravesicular sucrose and extravesicular glucose.

#### Preparation of oil-free GUVs using PVA gel-assisted hydration method

This protocol was adapted from Weinberger et al. (2013).[3] A 5% (w/w) solution of polyvinyl alcohol (PVA) was prepared by stirring PVA in water while heating at 90 °C. A volume of 10 µL of this solution was thinly spread into a well of a bottom-glass 96-well plate. The well plate was then placed in an oven at 70 °C for 45 min to dry the solution. A 1 g/L DOPC solution was prepared in dry chloroform. A volume of 3 µL of this solution was carefully spread on top of the dried PVA film, avoiding contact with the plastic walls of the well plate. The solvent was then dried under low pressure in a desiccator for 20 min or alternatively using a nitrogen flow. Next, 30 µL of inner solution (containing 1 µM Atto 565-labelled ssDNA reporter (a) in 1xTE buffer pH 8, 150 mM NaCl, 400 mM sucrose) was added onto the lipid film and left incubating for 35 minutes to allow gentle swelling and GUV formation. Afterwards, to detach the GUVs from the PVA gel, 100 µL of outer solution (1xTE buffer pH 8, 150 mM NaCl, 400 mM glucose) was added, and the whole solution was gently pipetted up and down several times. The whole solution of GUVs was then transferred to a clean microtube for storage or to another coated glass-bottom well for CLSM imaging.

#### Monitoring Chol-ssDNA flipping

To monitor DNA flipping, GUVs with a specific lipid composition (indicated for each experiment) were prepared, and a fluorescently labelled ssDNA reporter was added to the inner solution at a concentration of 1 µM during liposome preparation for its encapsulation. After GUV production, 50 µL of the GUV solution were placed in a well, and 0.3 µM of a complementary scavenger strand (NO cholesterol) was added and incubated for 5 minutes to bind any leaked DNA and avoid any false positives (main text Figure 3h-j). Following this, Chol-ssDNA was added at 0.5 µM (or at a different concentration when indicated). In experiments where glycerol is added to boost the yield of DNA flipping, glycerol was added to the GUV solution at the concentration indicated 5 minutes after Chol-ssDNA addition. An 80 µL volume of hexadecane was carefully layered on top of the aqueous solution to prevent sample evaporation. Imaging commenced immediately, as DNA flipping was observed in some GUVs within ~30 seconds after addition of Chol-ssDNA into the GUV solution. After each component addition into the GUV solution, the solution was gently pipetted to ensure proper mixing within the well.

#### Nuclease degradation assay

For the nuclease degradation assay, the flipping of Chol-ssDNA experiment was first performed as described above, using 1 µM of Cy5-labelled ssDNA reporter (b) encapsulated and 0.3 µM scavenger (b*) and 0.5 µM DY-530-labelled Chol-ssDNA (b*) added in the GUV solution. Next, DNase I was added to the GUV solution in the well plate to a final concentration of 1 g/L, followed by gentle pipetting for homogenization. The sample was imaged after 30 minutes and again after a 24-hour incubation at 4 °C.

#### $Ca^{2+}$ permeability experiment

GUVs composed of DOPC encapsulating 3 µM GECO and 1 µM Atto565-labelled ssDNA reporter (a) were prepared. After production, 50 µL of the GUV solution were placed in a well, followed by the addition of 0.3 µM ssDNA scavenger (a*) and 3 mM $CaCl_2$. After 5 minutes, Chol-ssDNA (a*) was added at a 0.5 µM final concentration, and the sample was imaged right after.

#### Dithionite assay

GUVs were prepared using DOPC and NBD-PE (0.1 mol%). The inner solution for encapsulation contained 1 µM of Alexa 647 and 1 µM of Atto565-labelled ssDNA reporter (a). After production, 50 µL of the GUV solution was placed in a pre-coated glass-bottom well, followed by the addition of 0.3 µM of ssDNA scavenger (a*). After 5 minutes, 0.5 µM of Chol-ssDNA (a*) was added in the GUV solution. A 100 mM stock of sodium dithionite solution was freshly prepared in 1 M Tris pH 10 before each experiment. To initiate NBD and Alexa 647 reduction, dithionite was added to the GUV solution on the well plate from the stock to a final concentration of 7.5 mM. Imaging was performed immediately after the addition. For single GUV fluorescence reduction traces, the GUV of interest was kept in focus and images were recorded every 1.7 s while imaging with both lasers simultaneously.

#### Fluorescence recovery after photobleaching (FRAP) experiments.

FRAP experiments were performed by bleaching a small circular region of interest (ROI) with a diameter of 2 µm for 2 seconds with a 100 % laser intensity. Post-bleaching images were recorded for 300 s. The intensities within the circular ROI were measured in ImageJ to quantify the recovery kinetics over time. The intensity data was fitted to a single exponential function:

$$F_{FRAP}(t) = A \times (1 - e^{-t/\tau})$$

where $\tau$ is the characteristic time constant. The half-life recovery time $t_{1/2}$ was extracted using the relation:

$$t_{\frac{1}{2}} = \ln 2 \times \tau$$

The diffusion coefficient was extracted following this equation:

$$D = \frac{\omega^2}{4 \times t_{\frac{1}{2}}}$$

where W is the radius of the bleaching spot (2 µm).

#### Transcription of RNA Spinach Aptamer upon DNA translocation

GUVs were prepared using DOPC. The inner buffer solution contained 40 mM HEPES, 125 mM KCl, 5 mM $MgCl_2$, and 400 mM sucrose, while the outer buffer was identical except that sucrose was replaced with glucose to create a density gradient. The inner solution was supplemented with the components required for *in vitro* RNA transcription: T7 RNA polymerase (1:15 dilution from the T7 RNA polymerase mix), 6.7 mM of each NTP (1:3 dilution from the NTP buffer mix), and 1 µM DNA template. Both the T7 RNA polymerase mix and the NTP buffer mix (containing 20 mM of each NTP) were sourced from the HiScribe T7 High Yield RNA Synthesis Kit. Additionally, the inner solution included 5 µM DFHBI—a fluorophore that binds the RNA spinach aptamer—and 0.3 µM Cy5.5-labeled ssDNA reporter (c), enabling membrane fluorescence feedback upon flipping. After GUV production, 0.1 µM ssDNA scavenger (c*) was added to the external solution, followed by 0.5 µM Chol-ssDNA (c*) five minutes later. To minimize evaporation, 80 µL of hexadecane was gently layered over the aqueous solution. Samples were incubated at 25 °C for 16 hours prior to CLSM imaging.

#### Image analysis

Images were analyzed using Fiji. Pre-processing steps included smoothing and despeckling to enhance image quality. For quantitative analysis, GUVs were imaged near the equatorial plane. Line profiles were obtained by drawing a straight line across the GUV and analyzing the gray values along this line. A line thickness of 20 pixels was chosen to average the intensity. To analyze the kinetics of DNA flipping, membrane fluorescence intensity over time was quantified by manually drawing a line along the fluorescent ring and saving it as a ROI. The gray values along this ROI were measured across the time series. The resulting intensity curve was then fitted to the same single exponential equation used for FRAP analysis (see above), enabling a comparison of the flipping kinetics with the diffusion coefficient of Chol-ssDNA anchored in the bilayer. This approach yields an apparent diffusion coefficient, that is limited by factors such as pore formation and thus does not strictly represent diffusion. To measure the intensity from the cavity, a circular ROI located within the lumen of the GUV was selected, and the gray value was measured across the time series. Background subtraction was performed by averaging the gray values from three areas without GUVs.

**Table S1. List of DNA oligonucleotides with their names, sequences, and the figures in which they were used.**

| | Name | Sequence (5’ → 3’) |
|---|---|---|
| **Figure 2-5** | ssDNA reporter (a) | GGA CGG CTC GGA TGC GC – **Atto 565** |
| | Chol-ssDNA (a*) | TCC GAG CCG TCC GCA TTC TAA TCA CC - **TEG-chol** |
| | ssDNA scavenger (a*) | GCG CAT CCG AGC CGT CC |
| | Scrambled Chol-ssDNA | TGA TTA TCG AGT TCA GTA TTT T - **TEG-chol** |
| | ssDNA without chol | TCC GAG CCG TCC GCA TTC TAA TCA CC |
| **Figure 2** | ssDNA reporter (b) | TGA TTA TCG AGT TCA GTA - **Cy5** |
| | Chol-ssDNA (b*) | **DY-530 -** TAC TGA ACT CGA TAA TCA AGT CTC ATA ATG GTT - **TEG-chol** |
| | ssDNA scavenger (b*) | TAC TGA ACT CGA TAA TCA |
| **Figure 5f** | 12-nt Chol-ssDNA | TCC GAG CCG TCC **-TEG-chol** |
| | 26-nt Chol-ssDNA | TCC GAG CCG TCC GCA TTC TAA TCA CC - **TEG-chol** |
| | 75-nt Chol-ssDNA | TCC GAG CCG TCC GCA TTC TAA TCA CCT TTT TTT TTT TTT TTT TTT TTT TT TTT TTT TTT TTT TTT TTT TTT TTT - **TEG-chol** |
| | ssDNA reporter | GGA CGG CTC GGA TGC GC – **Atto 565** |
| | ssDNA scavenger | GCG CAT CCG AGC CGT CC |
| **Figure 5g** | Sequence a | TCC GAG CCG TCC GCA TTC TAA TCA CC - **TEG-chol** |
| | ssDNA reporter | GGA CGG CTC GGA TGC GC – **Atto 565** |
| | ssDNA scavenger | GCG CAT CCG AGC CGT CC |
| | Sequence b | **DY-530 -** TAC TGA ACT CGA TAA TCA AGT CTC ATA ATG GTT - **TEG-chol** |
| | ssDNA reporter | TGA TTA TCG AGT TCA GTA - **Cy5** |
| | ssDNA scavenger | TAC TGA ACT CGA TAA TCA |
| | Sequence c | **Chol Prolinol** – TTT TTT TTT TTT CGC TAA TAC GAC TCA CTA TA |
| | ssDNA reporter | **Cy5.5 -** TAT AGT GAG TCG TAT TAG CG |
| | ssDNA scavenger | CGC TAA TAC GAC TCA CTA TA |
| **Figure 6** | ssDNA reporter (c) | **Cy5.5 -** TAT AGT GAG TCG TAT TAG CG |
| | Chol-ssDNA (c*) | **Chol Prolinol** – TTT TTT TTT TTT CGC TAA TAC GAC TCA CTA TA *(t7 promoter)* |
| | ssDNA scavenger (c*) | CGC TAA TAC GAC TCA CTA TA |
| | DNA template for RNA spinach aptamer | GGA GCT CAC ACT CTA CTC AAC AGT AGC GAA CTA CTG GAC CCG TCC TTC ACC CTA TAG TGA GTC GTA TTA GCG AGT ATA GGG |
| | Scrambled Chol-ssDNA | TGA TTA TCG AGT TCA GTA TTT T - **TEG-chol** |
| **Figure S4** | ssDNA reporter non-complementary to Chol-ssDNA b* | GGA CGG CTC GCA - **Cy5** |

Complementary DNA strands are marked with an asterisk (e.g., a and a*), indicating full or partial complementarity. Scavenger strands are fully complementary to ssDNA reporters. Underlined bases show the shared complementary domain between Chol-ssDNA and the ssDNA reporter.

### Supplementary Figures

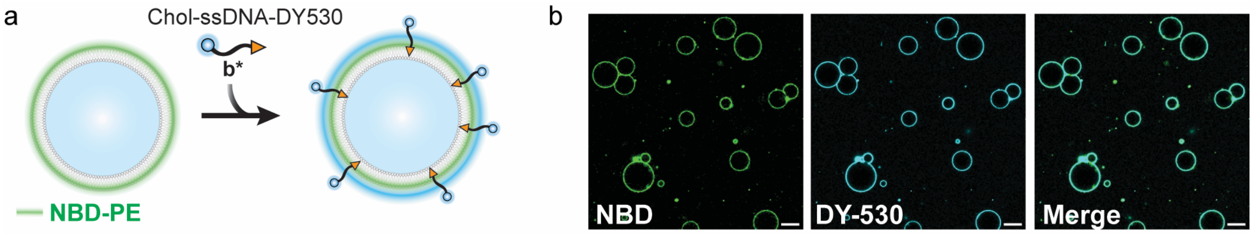

**Figure S1. Anchoring of Chol-modified ssDNA to all available GUV membranes.** Addition of DY-530-labelled Chol-ssDNA (b*) into a GUV solution labelled with NBD-PE results in the colocalization of the DNA on all the membranes. Conditions: DOPC + 0.1 mol% NBD-PE GUVs. Outer solution: 1xTE buffer pH 8, 150 mM NaCl, 400 mM glucose; 0.5 µM DY-530-labelled Chol-ssDNA (b*). Inner solution: 1xTE buffer pH 8, 150 mM NaCl, 400 mM sucrose. Scale bar: 20 µm.

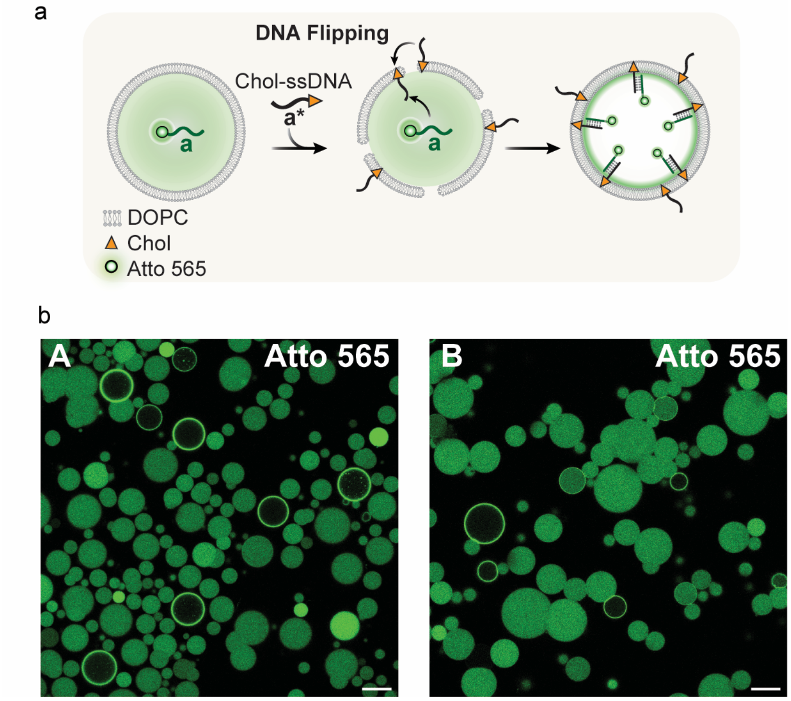

**Figure S2. Control experiments testing the influence of polyvinyl alcohol (PVA) on DNA flipping. a** Schematic representation of the Chol-ssDNA flipping experiment. **b** CLSM images after Chol-ssDNA flipping under different conditions: (**A**) BSA coating: The PVA coating on the well plates for imaging is replaced with a bovine serum albumin (BSA) coating. DNA flipping occurs to the same extent, indicating that any potential interactions between the GUV membrane and PVA do not affect the process. (**B**) BSA coating and free PVA: The PVA coating is replaced with a BSA coating as in (A), and 0.5% w/v PVA is added to the outer solution before adding Chol-ssDNA to assess whether PVA in solution influences membrane dynamics and therefore DNA flipping. After adding Chol-ssDNA, DNA flipping still occurs, suggesting that PVA has no significant effect. Conditions: DOPC GUVs. Outer solution: 1xTE buffer pH 8, 150 mM NaCl, 400 mM glucose. 0.3 µM ssDNA scavenger (a*); 0.5 µM Chol-ssDNA (a*). Inner solution: 1xTE buffer pH 8, 150 mM NaCl, 400 mM sucrose. 1 µM Atto 565-labelled ssDNA reporter (a). Scale bar: 20 µm.

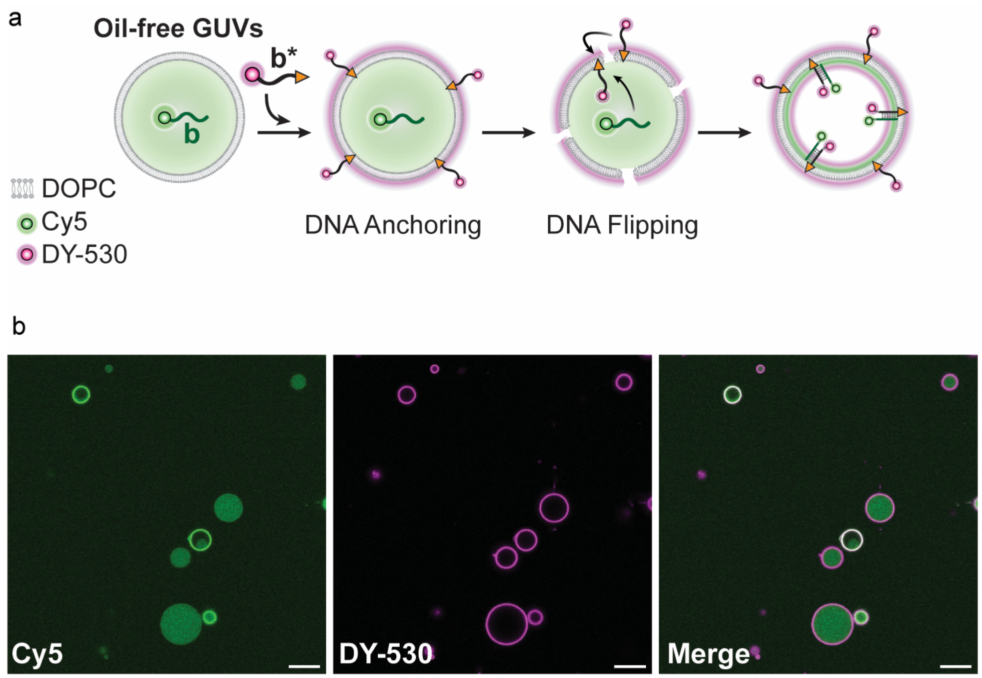

**Figure S3. DNA flipping in oil-free GUVs prepared via hydration.** To assess whether any potential residual oil in the membranes of GUVs produced by the emulsion transfer method affects DNA translocation, GUVs were also prepared using a hydration-based method[3] (see Experimental Section), which does not involve any oil. DNA translocation still occurs, indicating that any potential oil traces do not play any role in the process. Conditions: DOPC GUVs. Outer solution: 1xTE buffer pH 8, 150 mM NaCl, 400 mM glucose. 0.3 µM ssDNA scavenger (b*); 0.5 µM DY-530-labelled Chol-ssDNA (b*). Inner solution: 1xTE buffer pH 8, 150 mM NaCl, 400 mM sucrose; 1 µM Cy5-labelled ssDNA reporter (b). Scale bar: 20 µm.

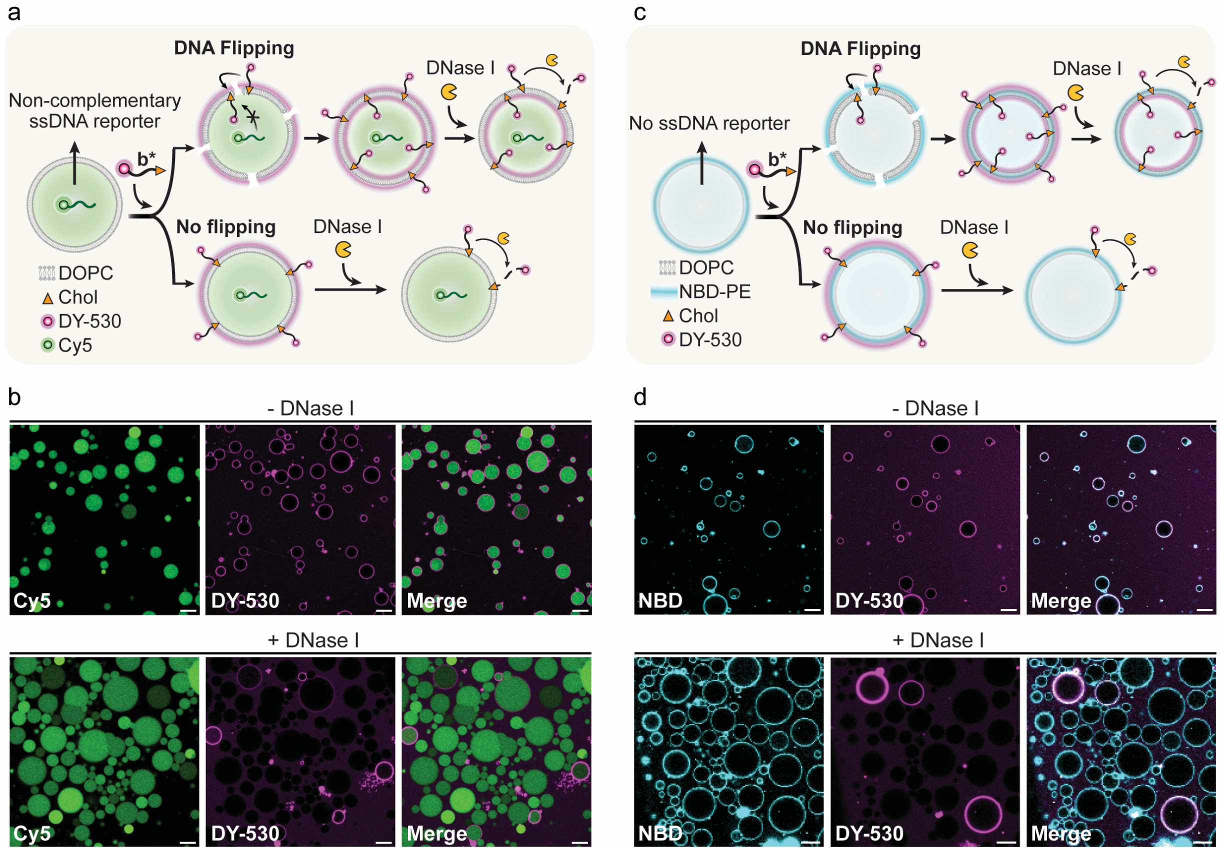

**Figure S4. Nuclease degradation assay in control GUVs to test the effect of encapsulated DNA on flipping. a** Schematic representation of GUVs that encapsulate a ssDNA that is non-complementary to Chol-ssDNA b*, as a control to test the influence of the encapsulated complementary strand for the DNA flipping. The DY-530 dye-labelled Chol-ssDNA b* enables tracking of the strand's location following nuclease degradation. **b** CLSM images before and after DNase I treatment showing that flipping events still occur, visible as membrane rings in the DY-530 channel in the presence of DNase I, regardless of the ssDNA sequence encapsulated. **c** A control experiment with no DNA encapsulated inside the GUVs to further assess the influence of encapsulated DNA on DNA flipping. **d** CLSM images before and after DNase I treatment showing that DNA flipping still occurs in GUVs with no DNA encapsulated, as evidenced by the presence of DY-530 fluorescent rings in the DNase I-treated GUVs. Conditions: DOPC GUVs (panel a, b). DOPC + 0.1 mol% NBD-PE GUVs (panel c, d). Outer solution: 1xTE buffer pH 8, 150 mM NaCl, 400 mM glucose; 0.5 µM DY-530-labelled Chol-ssDNA (b*); 1 g/L DNase I. Inner solution: 1xTE buffer pH 8, 150 mM NaCl, 400 mM sucrose; 1 µM non-complementary ssDNA reporter (panel a, b). Scale bar: 20 µm.

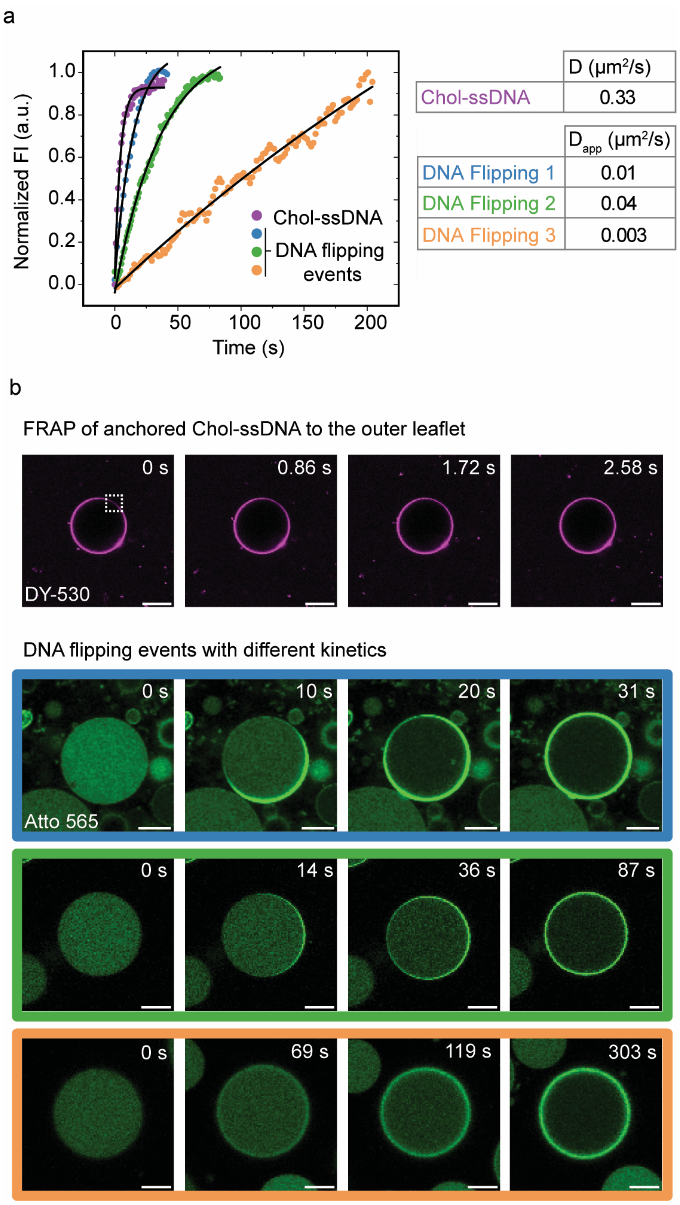

**Figure S5. Diffusion study of membrane-anchored Chol-ssDNA using fluorescence recovery after photobleaching (FRAP) and comparison of its diffusion rate with the kinetics of DNA flipping. a** Fluorescence recovery curve obtained by FRAP[4] for Chol-ssDNA anchored to the outer leaflet (magenta), plotted alongside three representative individual DNA flipping events (blue, green, orange), showcasing the emergence of membrane fluorescence (normalized) over time during the flipping process. Note that each DNA flipping event shows distinct kinetics. The events shown here represent the most characteristic examples selected for clarity and analysis. Diffusion (D) and apparent diffusion coefficients ($D_{app}$) were extracted by fitting the recovery curves for Chol-ssDNA, as well as the membrane fluorescence profiles during DNA flipping events, to a single-exponential function (solid line, see Methods). The results are summarized in the table. **b** Time-lapse CLSM images showing (top) FRAP of Chol-ssDNA anchored to the outer leaflet of a GUV and (bottom) time-lapse images of three distinct DNA flipping events with varying kinetics. The dashed square in b indicates the bleached area in the FRAP experiment. Conditions: DOPC GUVs. Outer solution: 1xTE buffer pH 8, 150 mM NaCl, 400 mM glucose; 0.5 µM DY-530-labelled Chol-ssDNA (b*) (panel a, b, FRAP); 0.3 µM ssDNA scavenger (a*) (panel a, b, DNA flipping events); 0.5 µM Chol-ssDNA (a*) (panel a, b, DNA flipping events). Inner solution: 1xTE buffer pH 8, 150 mM NaCl, 400 mM sucrose; 1 µM Atto565-labelled ssDNA reporter (a) (panel a, b, DNA flipping events). Scale bar: 10 µm.

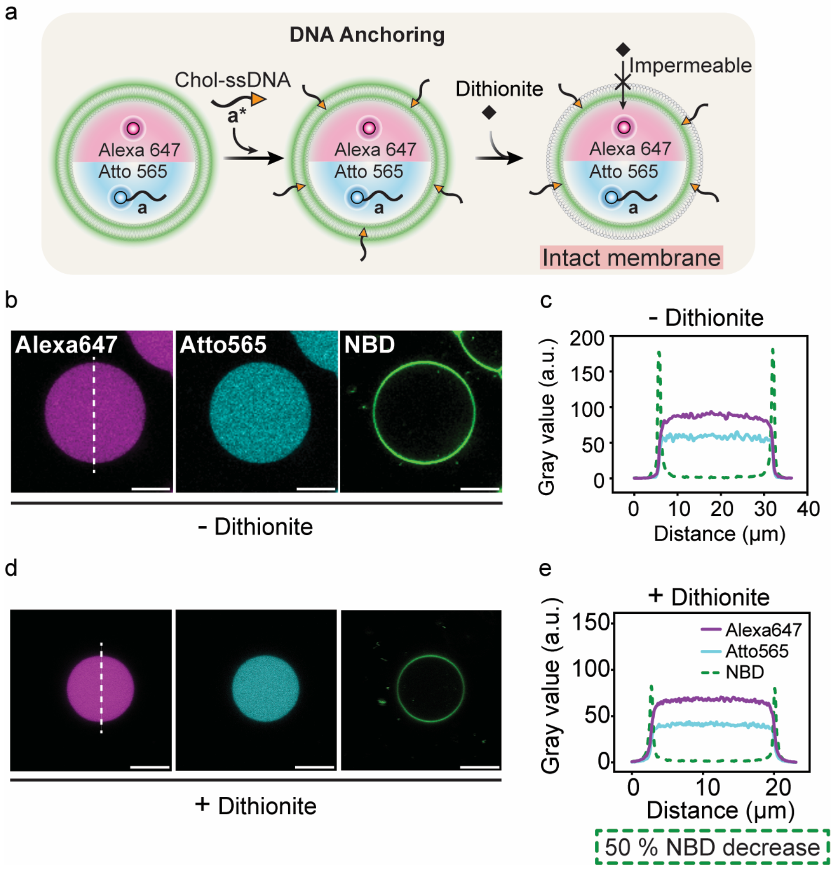

**Figure S6. Dithionite assay on GUVs with Chol-ssDNA anchored to the outer leaflet. a** Schematic representation of GUVs where Chol-ssDNA remains anchored without flipping. The membrane is expected to remain impermeable to dithionite, with no lipid flip-flop happening and therefore intact. **b** CLSM images of a GUV without DNA flipped (no ring in the Atto 565 channel) before dithionite reduction. Alexa 647 is encapsulated inside the GUV, while NBD is located in the lipid membrane. Signal changes after dithionite addition will probe membrane structure. **c** Line profile analysis of the CLSM images in b, prior to dithionite treatment. **d** CLSM images of a GUV without flipped DNA (no ring in the Atto 565 channel) after dithionite treatment. The Alexa 647 signal remains, while the NBD membrane fluorescence is reduced due to quenching of outer leaflet lipid exposed to dithionite. This partial quenching confirms that the membrane remains impermeable—and therefore intact—as complete quenching would suggest loss of membrane integrity. **e** Line profile analysis of the CLSM images in d, showing a ~50% decrease in NBD fluorescence. Conditions: DOPC + 0.1 mol% NBD-PE GUVs. Outer solution: 1xTE buffer pH 8, 150 mM NaCl, 400 mM glucose. 0.3 μM ssDNA scavenger (a*); 0.5 μM Chol-ssDNA (a*); 9 mM sodium dithionite. Inner solution: 1xTE buffer pH 8, 150 mM NaCl, 400 mM sucrose; 1 μM Atto565-labelled ssDNA reporter (a); 1 μM Alexa 647. Scale bar: 10 μm.

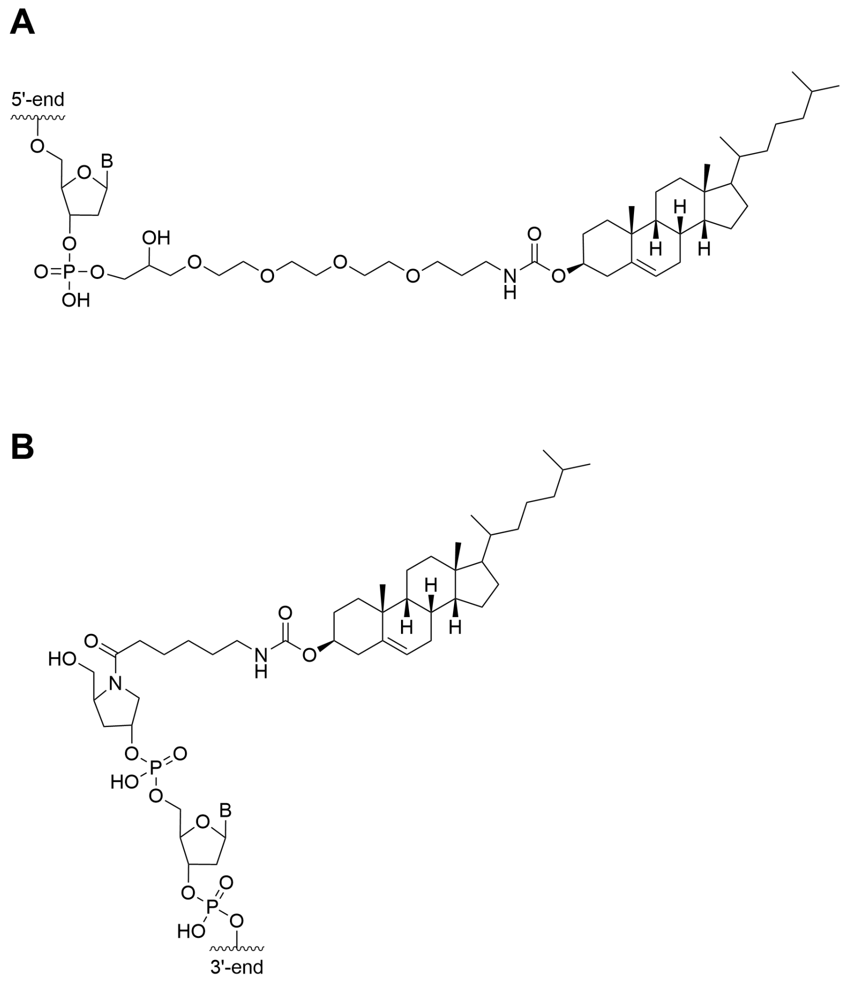

**Figure S7. Chemical structure of (A) 3'-Cholesterol-triethylene glycol (Chol-TEG) and (B) 5'-Cholesterol-prolinol.**

#### Supplementary Note 1. Theoretical calculation of the amount of Chol-ssDNA translocated across the membrane

To calculate the amount of Chol-ssDNA that flips across the membrane, we first calculate the number of encapsulated ssDNA reporter molecules per DOPC GUV. This number is given by:

$$N_{ssDNA,encapsulated} = C_{DNA} \times V_{GUV} \times N_A = C_{DNA} \times {}^{4}\!/_{3}\, \pi r^3 \times N_A = 1{,}84 \times 10^6\ molecules$$

where $C_{DNA}$ = 1 µM is the concentration of encapsulated ssDNA reporter, r = 9 µm is the average GUV radius, and $N_A$ is Avogadro's number.

After Chol-ssDNA flipping and hybridization to the ssDNA reporter, the reporter is completely depleted from the GUV cavity and localizes to the inner membrane surface. Since Chol-ssDNA and the ssDNA reporter hybridize in a 1:1 ratio to form Chol-dsDNA, the number of encapsulated ssDNA reporter molecules represents the **minimum** number of Chol-ssDNA molecules that have translocated across the membrane, $N^{min}_{Chol-ssDNA}$. Note that this assumption only holds because complete reporter depletion was observed.

$$N^{min}_{Chol-ssDNA} = N_{ssDNA,encapsulated}$$

The surface density of Chol-dsDNA on the inner leaflet of the membrane is then calculated by dividing the total number of molecules by the surface area of the GUV:

$$\rho_{Chol-dsDNA} = \frac{N^{min}_{Chol-ssDNA}}{4\pi r^2} = 1807\ {}^{molecules}\!/_{\mu m^2}$$

To assess the extent of membrane occupation, we estimate the membrane area available per Chol-dsDNA molecule:

$$A_{per\ Chol-dsDNA} = \frac{1}{\rho_{Chol-dsDNA}} = 5.54 \times 10^{-4}\ \mu m^2 = 554\ nm^2$$

To calculate the distance between molecules based on the area per Chol-dsDNA molecule, we can assume a hexagonal packing of the lipids:

$$A = \frac{3\sqrt{3}}{2} \times d^2$$

$$d_{Chol-dsDNA} = \sqrt{\frac{2 \times A_{per\ Chol-dsDNA}}{3\sqrt{3}}} = \sqrt{\frac{2 \times 554}{3\sqrt{3}}} = 14.6\ nm$$

Finally, the number of lipids on the inner leaflet is estimated using an area per DOPC lipid[5] of 0.7 nm$^2$:

$$N_{DOPC} = \frac{4\pi r^2}{A_{DOPC}} = 1.45 \times 10^9 \, molecules$$

The DNA-to-lipid ratio is defined as the ratio of the number of **Chol-ssDNA molecules**, $N_{Chol-ssDNA}^{min}$, to the number of **DOPC molecules**, $N_{DOPC}$, in the inner leaflet of the bilayer:

$$DNA \; to \; lipid \; ratio = \frac{N_{Chol-ssDNA}^{min}}{N_{DOPC}} = \frac{1}{787}$$

At a minimum, there is **1 Chol-dsDNA molecule** flipped into the bilayer for every **787 DOPC molecules** in the inner leaflet. The Chol-dsDNA have an average distance of 14.6 nm in a hexagonal lattice.

#### Supplementary Videos

**Supplementary Video 1**

Flipping is a defect-nucleated process

Flipping of Chol-ssDNA across the GUV membrane initiates at a localized point and rapidly propagates across the vesicle, resembling a defect-mediated process. The process is monitored through binding of fluorescent ssDNA reporter that hybridizes to the inner leaflet of the GUV membrane once the Chol-ssDNA flips to the inside.

**Supplementary Video 2**

Disturbances in the environment such as vesicle bursting can induce DNA flipping

Upon random bursting of a GUV, disturbances are generated in the surrounding environment, visibly affecting adjacent GUVs. One neighboring GUV responds by undergoing a clear DNA flipping event shortly after the initial disturbance. The first rectangle highlights the bursting GUV, while the second rectangle indicates the neighboring GUV undergoing DNA flipping.

**Supplementary Video 3**

DNA flipping affects membrane curvature

Membrane curvature changes can be inferred from the geometry of the contact interface between two interacting GUVs. Initially, the interacting surfaces are planar but following a DNA flipping event in one of the GUVs, a distinct change in geometry occurs, indicating a shift in membrane curvature. This suggests a potential link between DNA translocation and membrane curvature changes. Notably, Chol-ssDNA anchored solely in the outer leaflet is known to induce membrane curvature,[6] supporting the idea that its redistribution among leaflets during flipping could drive the observed geometric transitions.

**Supplementary Video 4**

Glycerol induces fusion between GUVs

The addition of 1%(v/v) glycerol to a GUV solution increases membrane dynamics, evidenced by the observed fusion events. Such events are not detected when glycerol is not present.
